# Stable twin rDNA loci form a single nucleolus in brewer’s yeast

**DOI:** 10.1101/2022.06.14.496147

**Authors:** Luciana Lazar-Stefanita, Jingchuan Luo, Max A. B. Haase, Weimin Zhang, Jef D. Boeke

## Abstract

The nucleolus is the most prominent membraneless compartment within the nucleus^1, 2^ - dedicated to the metabolism of ribosomal RNA. Nucleoli are composed of hundreds of ribosomal DNA (rDNA) repeated genes that form large chromosomal clusters^3–5^, whose high recombination rates can cause nucleolar dysfunction and promote genome instability^6–8^ related to metabolic and genetic diseases^9–13^. Intriguingly, the evolving architecture of genomes appears to have favored two strategic rDNA locations in a broad range of species – where a single locus per chromosome is situated either near the centromere or the telomere^14, 15^. To delve into how organisms may benefit from these nuclear organizations, we used a fused-karyotype strain of *Saccharomyces cerevisiae*^16^ to megabase-engineer a chromosome with twin chromosome-collinear rDNA loci. We showed that the twin-rDNA yeast readily adapts exhibiting wild-type growth and maintaining rRNA homeostasis. Using imaging and chromosome conformation capture, we found that the twin loci merge into a single subnuclear compartment throughout the cell cycle. Unexpectedly, we found that rDNA locus size is dependent on its position relative to the centromere, whereby the locus that is centromere–distal undergoes size reduction at a higher frequency compared to the centromere-proximal counterpart. In sum, our work sheds light on the structural evolution of rDNA loci and provides new tools to study the rDNA dosage effect on cellular metabolism.

## Introduction

The most abundant genes in cells are those encoding ribosomal RNA (rRNA), usually arranged as a series of tandem repeats that include transcriptional units for rRNAs spaced by non-transcribed sequences^3, 17^. As rRNAs are the structural components of ribosomes, they represent the most abundant RNA molecules in most organisms, accounting for 60-70% of cellular transcripts e.g., 10^5^/yeast cell^18^ and 10^6–7^/animal cell^1^.

In eukaryotes, rRNA transcription and ribosome assembly take place in a membraneless organelle that occupies a substantial proportion of the cell nucleus, namely the nucleolus^2, 19^. Ostensibly incongruous features of the nucleolus – such as rapid molecular turnover and liquid-like dynamics, yet maintaining a coherent shape – are reconciled by the liquid-liquid phase separation model^1, 20^, which proposes that rRNA and associated proteins self-organize based on their intrinsic biophysical properties^21^. Moreover, the structure and activity of the nucleolus are highly dynamic throughout cell division, as is evident in eukaryotes with open mitosis in which nucleolar breakdown and transcriptional shutdown are concomitant with entry into mitosis^22, 23^. Nucleoli assemble around distinct chromosomal clusters/loci, gathering hundreds of tandemly repeated rDNA genes that largely exceed the number of units needed for survival. Indeed, typically, only ∼50% of the rDNA genes are actively transcribed while the remainder are kept inactive, an equilibrium that can be readily tuned by altering the balance between euchromatin and heterochromatin status at this locus^24, 25^. Furthermore, a high rRNA level - positively correlated with growth rate in many microbes^26^ - can be achieved not only by boosting transcription but also by rDNA copy number expansion^27^. Given their repetitive nature and the high demand for rRNA transcripts, the rDNA loci are among the most unstable genome structures^7, 28^. Their instability relies on a repeat-mediated homologous recombination (HR) mechanism that requires both rDNA replication^27, 29^ and transcriptional processes^30, 31^ that often result in clonal size variation in the repeat number.

Cyto-taxonomic studies, based on the widespread usage of fluorescence in situ hybridization, enabled mapping of rDNA loci on chromosomes of thousands of species. Number and position of these loci, each containing hundreds of tandem repeats, were compiled and used for in silico analyses of their evolution in plants^14^ and animals^15^. To reveal that most taxa have genomes with multiple rDNA loci, with an average number of only three to four loci per diploid genome, indicating that most organisms tend to maintain a moderately low number of loci. In plants, locus number was positively correlated with genome size; whereas in animals no such correlation was observed. Surprisingly, although rDNA loci were identified at nearly any chromosomal position, they prevalently appear either pericentromeric and/or peritelomeric (in >75% of karyotypes). Genome-wide studies in mammals^32^ and yeasts^33^ provided evidence to support the hypothesis that rDNA arrays have changed their chromosomal position multiple times during evolution: in almost every case, the array disappeared entirely from the ancestral site, leaving behind a small ‘intergenic scar’^33^.

These insights hint at two potential aspects of rDNA evolution: (*i*) a common ancestral mechanism through which this locus may migrate within genomes and (*ii*) deleterious effects associated with more than a single rDNA locus per chromosome. Regarding the first hypothesis, we can only speculate that HR and ‘nonreciprocal crossover’ between repeats^34–36^ may be responsible for rDNA dynamics. Here we address the second hypothesis by monitoring the effects of a rDNA migration event in which the entire native megabase rDNA array was transplanted to an ectopic intra-chromosomal location through a copy-and-paste mechanism.

## Results

In budding yeast, such as *Saccharomyces cerevisiae*, the *RDN1* locus is approximately 1.5 Mb region that contains 150 – 200 copies of a 9.1 kb unit and accounts for ∼10% of the entire yeast genome^37^. Each repeat encodes four genes: 5.8S, 25S, 18S and 5S rRNAs transcribed by RNA polymerase I and III^38, 39^. The entire array exists in single copy on the right arm of chromosome *12*, located roughly 300 kb from *CEN12* and ∼600 kb from *TEL12R* ^4, 40^, and forms a single crescent shaped nucleolus apposed to the nuclear envelope^41^ (Figure 1a). To assess whether multiple rDNA loci can support yeast viability, previously constructed strains with a single locus on different chromosomes (chr *12* and chr *3*)^42^ were mated and the resulting diploid was sporulated. This allowed the isolation of a haploid strain carrying two rDNA clusters on heterologous chromosomes, validated through pulsed-field gel electrophoresis (PFGE) (Extended Data Figure 1a). Notably, this strain displays a single nucleolar compartment (Figure 1b; Extended Data Figure 1b). These results led us to conclude that yeast (just like many other eukaryotes) can readily adapt to live with multiple rDNA loci on different chromosomes. However, it leaves unanswered the question of whether the yeast genome can be engineered with multi-megabase rDNA loci coexisting on the same chromosome.

**Figure 1.**
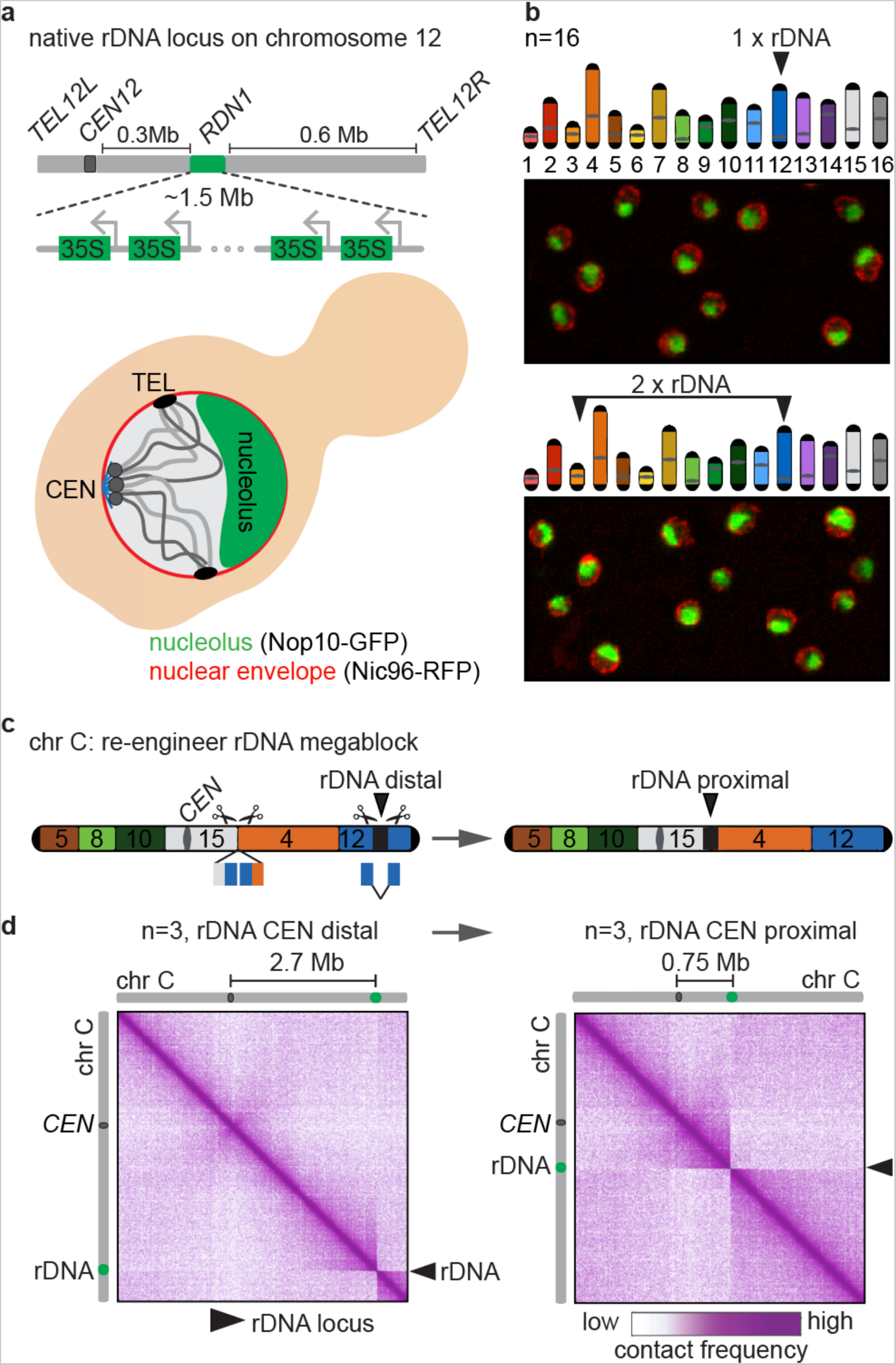
Transplantation of the rDNA megablock to ectopic genome locations. **a** Diagram showing wild-type chromosome *12* that contains the rDNA locus (green) and a simplified representation of nuclear organization in the wild-type yeast, where examples of chromosome arms (gray lines) are anchored at the nuclear membrane through centromeres (*CEN*) and telomeres (*TEL*). **b** Representative microscopy images of yeast nuclei with one or two rDNA loci (black arrowheads in the schematic indicate rDNA containing chromosomes). **c** Schema illustrating the relocation (arrowheads) of the rDNA megablock (black box) on the fused megachromosome *C* (chr *C*: color code corresponds to the design of chromosome fusion) in yeasts with *n*=3 chromosomes. Colored blocks indicate sequence homology of the repair donors at CRISPR-Cas9 cutting sites. **d** Hi-C contact maps of chr *C* shown in gray on the *x*- and *y*-axis of each map. The rDNA locus is *CEN* distal on the left, whereas, after transplantation it appears *CEN* proximal on the right. rDNA distance from the centromere (dark gray) is indicated atop the maps.

### rDNA Megablock engineering: one step transplantation of a megabase locus to a target position

To answer the above question, we used a karyotype-engineered yeast strain carrying three megabase-sized chromosomes (*n*=3; with chromosomes *A*, *B* and *C*) that have been previously generated through sequential telomere fusion events of the sixteen wild-type chromosomes^16^. These genome reengineering steps gave rise to megachromosome *C* (5.5 Mb-long), that resulted in a dramatic change in the position of the rDNA locus relative to the nearest active centromere (left drawing in Figure 1c, the black arrowhead points to the rDNA distal position). Precisely stated, on chr *C,* the rDNA-centromere distance increased 9-fold relative to the wild-type distance, reaching up to ∼2.7 Mb, whereas its native position relative to the nearest (right arm) telomere remained unchanged. Not only did this ‘passive’ migration of the locus to a distal location not affect growth (see JL411 strain generated by Luo et al.^16^), but it also provided a chromosomal arm long enough to allow for an insertion of a second centromere-proximal locus in *cis* (right drawing in Figure 1c). Here we developed a variation of the CRISPR-Cas9 editing approach to engineer megachromosomes that we call ‘Megablock engineering’ (Figure 1c; Methods), in which a megabase-sized segment of a chromosome is moved to a target location, potentially anywhere in the genome. Megablock engineering deploys two CRISPR guide RNAs that cut at the boundaries of the genomic DNA ‘block’ to be relocated, and a third guide cutting at the target location. Additionally, three ‘linker DNAs’ are provided: one that joins the native rDNA locus flanking ends together, and two that join the left and right junctions of the target locus to the ends of the megablock (Figure 1c). In this case, the entire megabase long rDNA locus was removed from its native centromere-distal location and relocated 2 Mb closer to the chr *C* centromere. This extremely aggressive genome engineering maneuver was surprisingly efficient, with 6/41 colonies score by PCR genotyping showing evidence of successful megablock engineering. Hi-C was used to structurally validate the rDNA locus transplantation (Figure 1d, black arrowheads point at the rDNA boundaries). One such strain (JL665), with *CEN* proximal rDNA was then used for further analyses and experiments. Of note, the orientation of the transplanted rDNA array relative to the chromosome was maintained to ensure locus directionality (e.g., rDNA replication and transcription orientation occur telomere-to-centromere in both the JL665 strain and its JL411 parent with *CEN* proximal or distal rDNA loci, respectively). By mating ‘parental’ and ‘transplanted’ strains, with either centromere-distal or centromere-proximal rDNA, respectively, we generated a heterozygous diploid with two loci asymmetrically positioned on each homolog of megachromosome *C*. Finally, haploids with duplicated rDNA loci in *cis* were isolated by sporulation of this diploid strain and their integration junctions were PCR-verified (Extended Data Figure 1c, d).

### Effects of rDNA duplication on fitness

Whole genome sequencing validated the duplication of the rDNA locus in the haploid strains (LS71 with ‘twin’ rDNA loci) by estimating twice as many rDNA-mapping reads in 2xrDNA/chr *C* (Figure 2a). Importantly, the newly formed rDNA junction sequences were confirmed and no chromosomal abnormalities potentially caused by recombination events between the two collinear loci were detected (Extended Data Figure 2a). Not only did the 2xrDNA isolates lack obvious fitness defects (Figure 2b; Extended Data Figure 2b), but they rapidly adapted to the excess of rDNA repeats by fine-tuning the rRNA levels (Figure 2c; Extended Data Figure 2c). In addition, a small number of genes were differentially expressed in the 2xrDNA vs. 1xrDNA strains (Figure 2d; Extended Data Table 2) and represented a nonrandom coherent set of genes. Notably, genes involved in the metabolism of glucose and metal transport were downregulated, whereas phosphate starvation-induced genes were upregulated. These results suggest that twin-rDNA cells may experience a delayed metabolic transition from glucose fermentation (fast-growth) to respiration (slow-growth) in response to nutrient deprivation.

**Figure 2.**
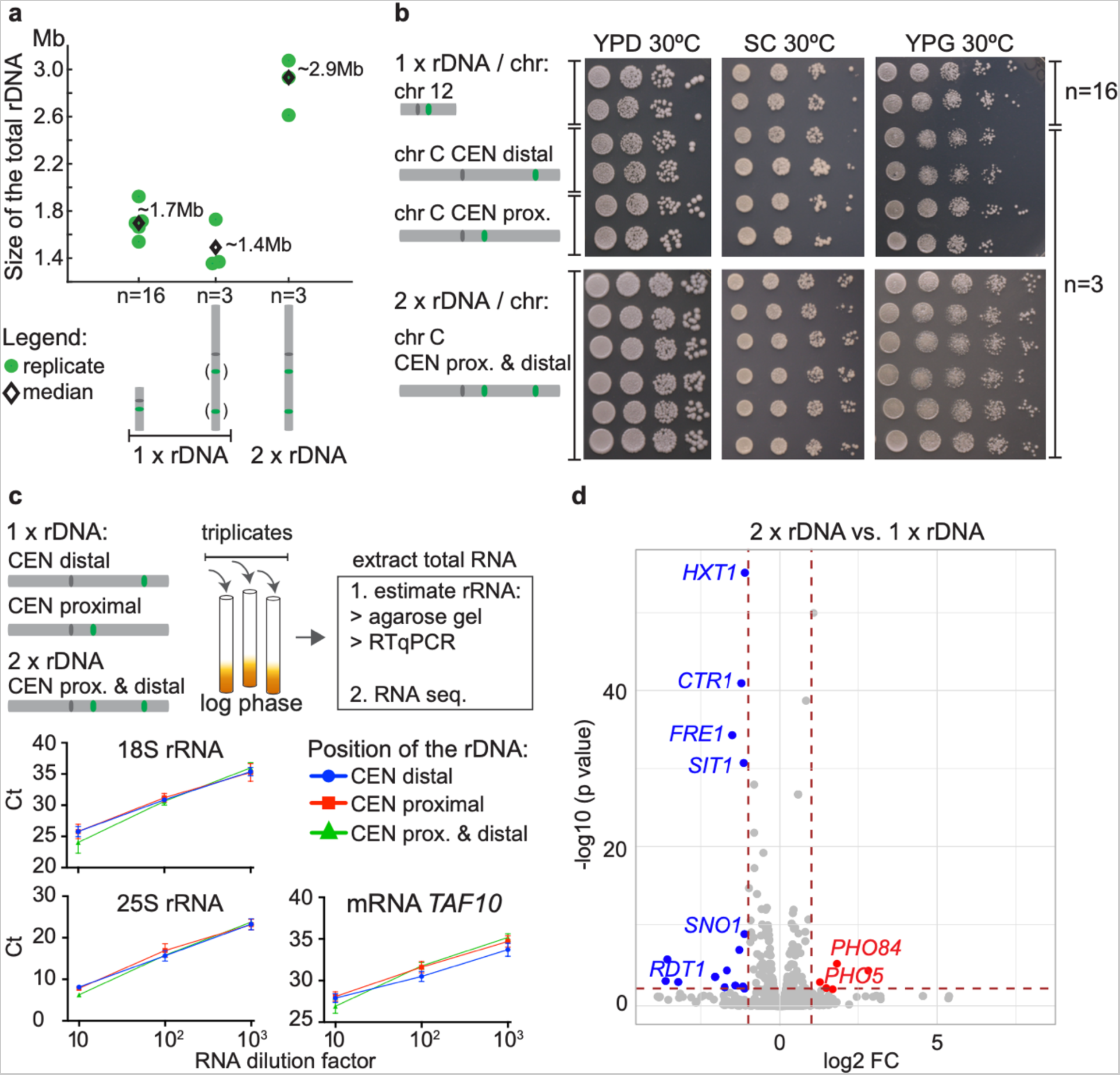
Effects of rDNA duplication on cell fitness. **a** Estimate the total length of the rDNA by deep sequencing: % of rDNA aligned reads normalized by genome size (12.3 Mb). Megachromosome *C* schematics are shown on the bottom of the plot (gray), where round brackets indicate either the proximal or the distal rDNA locus (green). **b** Serial dilution assay on different growth conditions at 30°C of *n*=16 and *n*=3 isolates with either single or duplicated rDNA. **c** Schematics of RNA-based assays in *n*=3 isolates with 1xrDNA vs 2xrDNA. RTqPCR plots on three serial (1:10) dilutions of total RNA are used to estimate changes in the amount of 18S and 25S ribosomal RNAs relative to the control messenger RNA, *TAF10*. **d** Volcano plot showing RNA-seq data comparing transcriptomes of 1xrDNA and 2xrDNA strains in triplicates. Red and blue dots indicate differentially expressed genes in 2xrDNA vs. 1xrDNA: in blue downregulated in red upregulated (*P* < 10−5, |fold change| > 2).

Links between function and structure of the rDNA locus in yeast have been previously described, and showed that a subset of the repeats are usually actively transcribed while the remainder are silenced^25, 43^. Their transcription level was correlated with distinct chromatin states, defined by the association of the repeats with DNA-folding markers of either euchromatin (e.g., High-mobility group proteins, Hmo1^44^) or heterochromatin (e.g., silencing complex, SIR^45^). Here we identified mild changes in the expression level of these chromatin modifiers: the rDNA-silencing factor (Sir2) was upregulated, whereas the active rDNA copy-associated protein Hmo1 was downregulated in 2xrDNA compared to 1xrDNA strains (Extended Data Figure 2d). This transcriptional anticorrelation is consistent with an excess of repeats in twin-rDNA strains that may demand additional silencing.

Next, we investigated the structural organization of chr *C* with twin-rDNA loci in *cis,* and their chromosomal (in)stability.

### Structure of chromosomes with twin-rDNA loci in cis

We used Hi-C^46, 47^ to probe the 3-dimensional (3D) structure of genomes with twin-rDNA arrays. The 2D contact maps of chr *C* containing a single locus, either centromere-proximal or distal, were combined in a single control plot. For comparison purposes, the control plot is displayed adjacent to that obtained from strains with *cis* twin-rDNA loci (Figure 3a). In the control plot, two indistinguishable boundaries formed at each rDNA location, as a result of similar insulation rates from their corresponding adjacent regions. In contrast, the two boundaries in *cis* appeared to give rise to distinct contact frequencies, where the insulation (‘sharpness’) at the *CEN* proximal rDNA cluster prevailed over the distal twin. This observation was validated through contact quantification plots for each locus that considered contacts between the intervening region enclosed by the two loci and the corresponding non-intervening rDNA-flanking regions (highlighted by blue dashed rectangles in Figure 3a). From this analysis we observed that the distal locus forms a weaker chromosome boundary (lower degree of insulation) when its centromere-proximal twin is located in *cis* (Figure 3b), a result that suggested distinct levels of structural ‘bulkiness’ between the *cis* rDNA loci.

**Figure 3.**
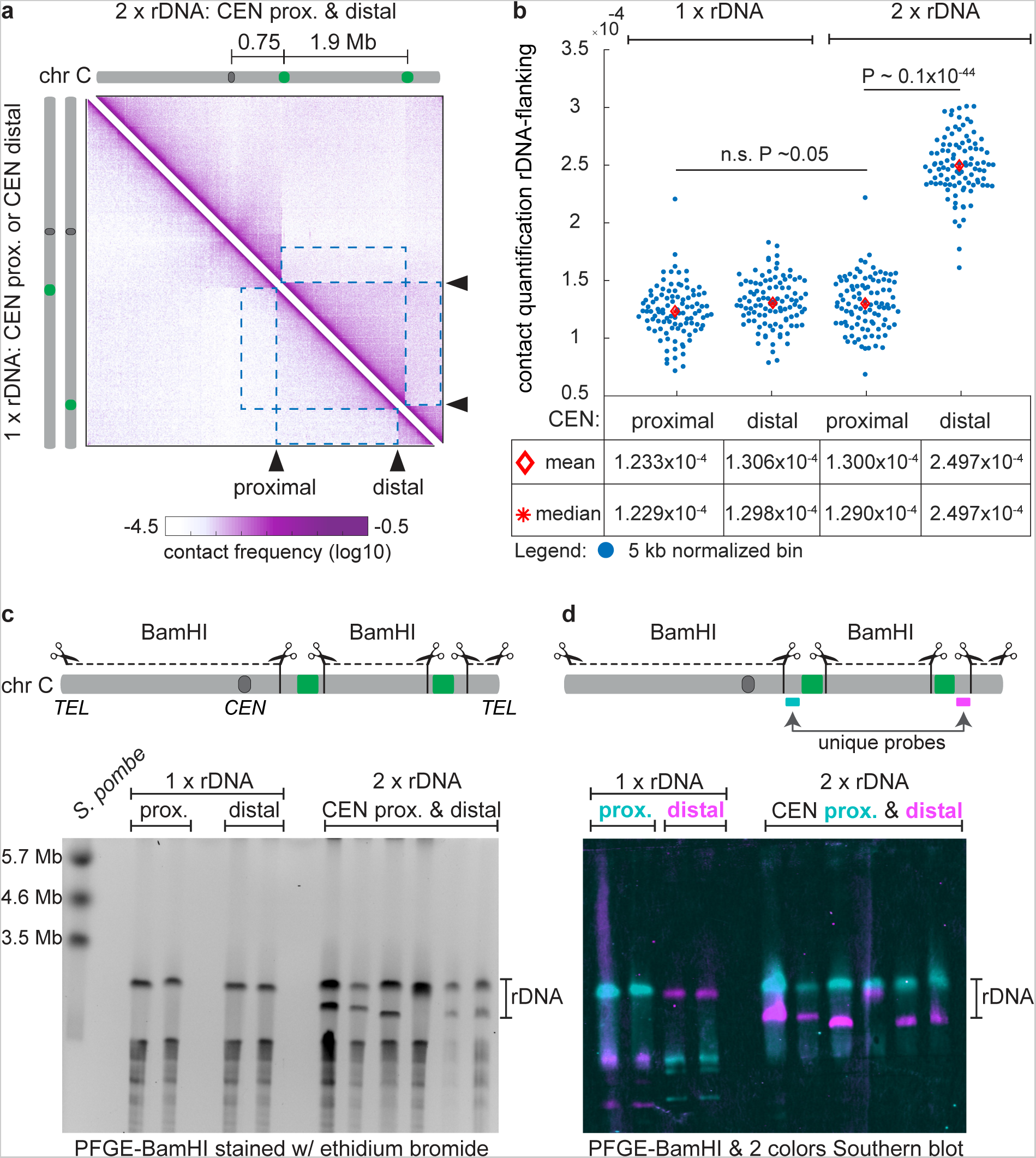
Stability of the rDNA locus is a function of chromosome position. **a** Hi-C contact maps of megachromsome *C* (chr *C* in gray on the *x*- and *y*-axis of each contact map) in *n*=3 strains with either one or two rDNA loci. Bottom left triangle map: average representation of two merged contact maps with 1xrDNA per chr *C*, either *CEN* proximal or *CEN* distal. Top right map: average representation of three merged contact maps with 2xrDNA per chr *C*, *CEN* proximal and distal. rDNA distance from the centromere (dark gray) is indicated atop the map. **b** Contact quantification between rDNA-flanking regions on chr *C*, highlighted in blue rectangles in panel (a): 500 kb sequence, either upstream *CEN* proximal or downstream *CEN* distal, and the 1.9 Mb sequence, downstream or upstream, respectively. Mean, median of absolute contact values and their *P* values are indicated for each strain with either one or two rDNA loci. **C** - **d** Estimate size of the rDNA arrays. PFGE of BamHI digested chromosomes (c) and the corresponding Southern blot (d) with unique probes designed for each rDNA position. Schematic indicates the position of rDNA arrays and probes (cyan or fuchsia) on chr *C*. PFGE run specifications: *S. pombe* program for mega-size chromosome separation.

Given that the 2xrDNA map represents consistent results obtained from 3 independent yeast isolates, we reasoned that the structural difference between the twin loci may arise from distinct epigenetic or genetic effects. Two major chromatin modifications at the distal locus could potentially be responsible for this divergence: (*i*) increased heterochromatin that may result in a highly condensed/compacted structure, and/or (*ii*) a reduction of rDNA repeat copy number. As the two loci are identical in sequence, we were unable to distinguish between rRNA transcripts originating from either locus (see discussion). Even though we may be unable to rule out the hypothesis of different levels of heterochromatin at the two *cis* rDNAs, their overall lengths can be directly measured. The size of each rDNA locus was estimated using PFGE and dual color Southern blotting on BamHI-digested chromosomes (Figure 3c, d; Methods). After digestion with BamHI, which does not cut within the rDNA repeat unit, the rDNA loci were left with unique DNA-flanking sequences, utilized to design probes that specifically recognize each locus. Surprisingly, all 2xrDNA isolates showed a consistent and locus-specific size reduction of the distal rDNA locus of ∼500-700 kb when compared to the 1xrDNA-distal parent strain (Figure 3d; Extended Data Figure 3a, b). Also, a higher clonal variability in the size of the distal locus was observed in isolates passaged for ∼100 generations (Extended Data Figure 3c). These results led us to hypothesize that rDNA repeats in yeast are more stable when located centromere-proximal rather than distal. To our knowledge this is the first evidence of a direct correlation between size of a repeated region and its relative genome location.

Finally, we probed the stability of the two loci in *cis* in function of Sir2, known to suppress recombination between repeated ribosomal genes^30, 45^. We found that the two distinct rDNA bands were lost (replaced by DNA smear) in all Δ*sir2* isolates (Extended Data Figure 3d), supporting the idea that the specific size variation of the two loci in *cis* may depend on differential Sir2 activity.

### Mitotic reorganization of the nucleolus with cis rDNAs

For the nucleolus to be correctly segregated between mother and daughter cells in late anaphase, rRNA transcription needs to be shut down to achieve rDNA-cluster condensation^48, 49^. Importantly, these condensation processes mainly involve the rDNA and are less pronounced across the remainder of the yeast genome.

Our previous work showed that the mitotic reorganization of chromatin is preserved along mega-chromosomes (Lazar-Stefanita et al., 2022 in press). Here, we specifically addressed the structure of megachromosome *C* with twin *cis* rDNAs and that of the resulting nucleolus during the cell cycle. Overall we expected chr *C* to undergo a typical mitotic reorganization^50^, however it remained unknown how the ∼1.9 Mb-long non-rDNA sequence linking the two loci may interfere with the structure of the nucleolus. Cells synchronized in G1 (alpha-factor) and in anaphase (*cdc15-2* ts^51^) were imaged and showed very clearly that the two rDNAs consistently formed a single nucleolar mass per cell throughout mitosis (Figure 4a; Extended Data Figure 4a).

**Figure 4.**
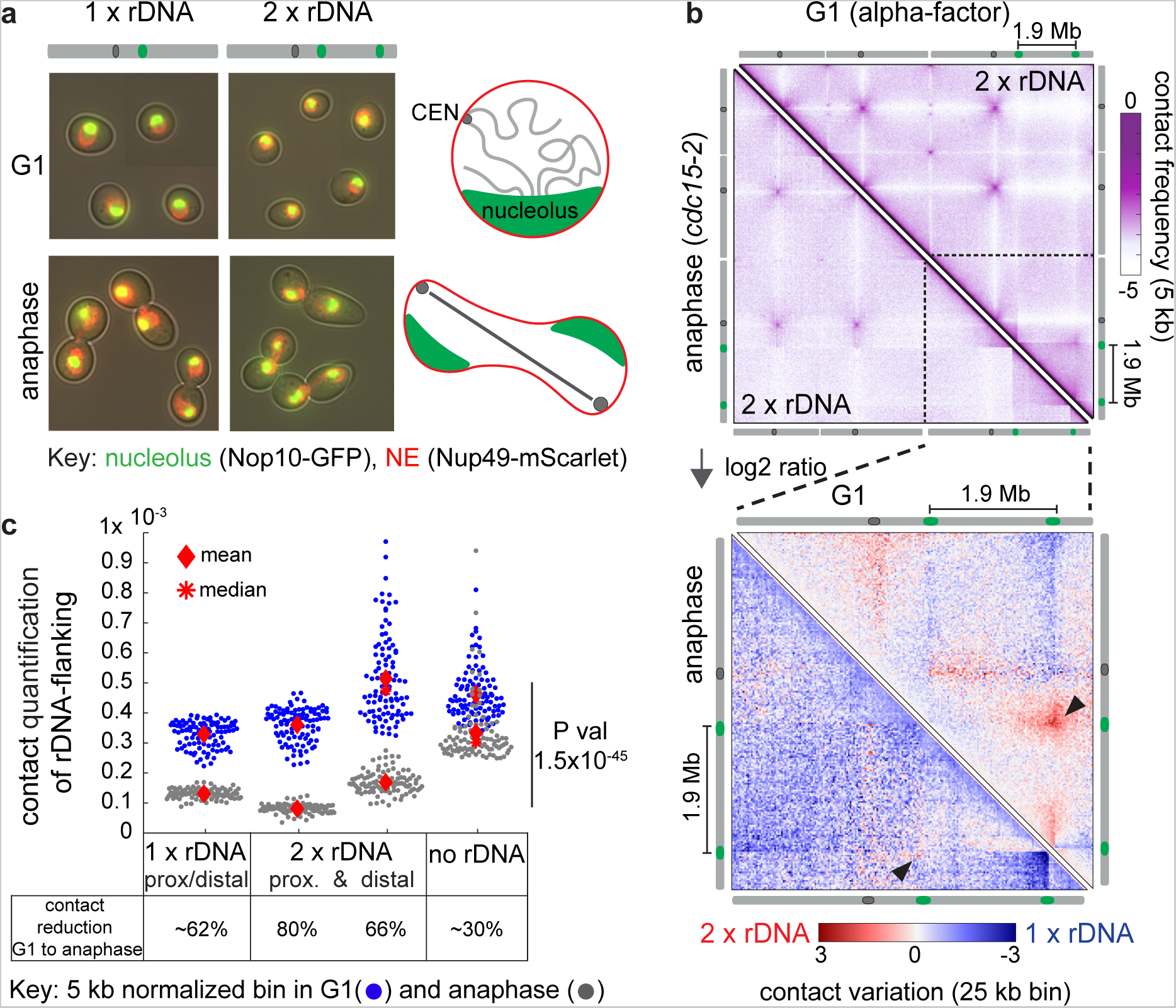
Twin rDNA loci in *cis* form a single nucleolus during cell division. **a** Representative microscopy images of *n*=3 strains with one or two rDNA loci in *cis*. Cells were synchronized in G1 (alpha-factor) and late anaphase (*cdc15-2* ts) before imaging. **b** Top panel: Hi-C maps of megachromsomes in *n*=3, with two rDNA loci in *cis* on chr *C* (dashed outline), from cells synchronized in G1 (top right map) and late anaphase (bottom left map). Bottom panel: contact variation maps (log2-ratio) of chr *C* with 1xrDNA vs. 2xrDNA in G1 (top right map) vs. late anaphase (bottom left map). Black arrowheads point at contact variations adjacent to the rDNA loci. **c** Contact quantification between rDNA-flanking regions on chr *C*, as described in Figure 3b. Relative percentages of contact reduction in anaphase compared to G1 are indicated for each strain with one, two and no rDNA.

The Hi-C maps revealed yet another layer of structural complexity of the nucleolus. The G1 map displayed strong contacts between the two rDNAs in *cis*, but these were lost in late anaphase (Figure 4b, black arrowhead; Extended Data Figure 4c). Contact quantifications between the rDNA-flanking regions (as described in Figure 3a, b) showed an overall decrease in anaphase compared to G1 (Figure 4b), in agreement with their increased condensation during anaphase that may strengthen chromosome boundaries. Note that the folding status of the rDNA-intervening sequence on chr *C* is not influenced by the rDNA-flanking loci (Extended Data Figure 4b, contact probability *p*(*s*) decay). Therefore, we conclude that (*i*) the ∼1.9 Mb rDNA-intervening sequence does not interfere with the organization of the nucleolus, and that (*ii*) the two loci are treated as distinct chromosomal entities during anaphase condensation.

## Conclusion and discussion

Here we provided the first living evidence of a eukaryotic genome carrying more than one rDNA locus located within the same chromosomal arm. The remarkable stability of the yeast genome is highlighted in its transcriptomics and the homeostatic control of the rDNA repeats at each locus. The structure of the twin-rDNA nucleolus and its reorganization during cell cycle was investigated using microscopy, showing a high degree of structural conformity between 1xrDNA and 2xrDNA strains.

### Design of the strains

Prior works on rDNA have employed interspecies hybrids to address if any differential stability of this locus in different yeasts^52–54^. Most of these studies observed a uniparental loss of rDNA genes, a relevant example is the hybrid between *S. cerevisiae* and *S. eubayanus* in which the latter’s locus collapsed completely whereas the *S. cerevisiae* remains full sized. Therefore, we reasoned that only by using two native and sequence-identical rDNA arrays located at different positions along the chromosome we could truly evaluate the position-effect on their structure and stability.

The ectopic location for rDNA mega-transplantation corresponds to a chromosome fusion site, specifically the right and left ends of chr *15* and chr *4*, respectively. Given that these regions were initially subtelomeric, which typically lack essential genes, we reasoned that they would be ‘insensitive-to-insertion’ sites for the huge rDNA locus. Finally, the two loci in *cis* conserved the wild-type transcriptional orientation (telomere-to-centromere). Our results show a consistent reduction in size of the centromere distal locus compared to the proximal counterpart. This peculiar size readjustment appears to be Sir2-dependent (Extended Data Figure 3d) suggesting that the transcriptional activity at the two loci maybe different. We speculate that the centromere-distal locus, which is flanked by its native sequence, is predominantly transcribed and therefore more prone to rapid size adjustments. Thus, explaining the homeostasis in rRNA level we observed.

### Nucleolus as a liquid condensate

We found that the yeast rDNA readily forms a single nucleolus, independent of its: position in the genome (homologous or heterologous chromosomes), number of loci per chromosome (single or duplicated) and repeat size. This is in full agreement with the proposed liquid-liquid phase separation model for the nucleolus^1, 21^, which envisions the intrinsic biochemical properties of the rDNA as major driving forces for nucleolar assembly that can act over very long genomic distances.

### Potential consequences of a twin-collinear organization of the rDNA loci

Evolutionary analyses of rDNA loci suggested that selective pressure may act to preserve specific characteristics (such as structure, number, and position) of these highly repeated loci in eukaryotes. Here we found that placing two rDNA loci in *cis* increases the rate of abnormal meiotic recombination (Extended Data Figure 5), possibly due to mispairing of the homologous chromosomes driven by the rDNA loci. We speculate that the misaligned chromatids can undergo unequal crossing-over resulting in an increased frequency of nonreciprocal exchange of genetic information (or gene conversion) and concomitant spore lethality. Our current model suggests that a high meiotic fitness cost is associated with multi-collinear repeated loci, and this would be predicted to ultimately be purged from a sexual population.

In addition, the extra copy of the rDNA locus appears to affect the metabolic response to nutrient consumption that would normally induce cells to exit the fast-growing phase, enter slow-growth and eventually arrest. Our transcriptomics data seem to suggest that the rDNA surplus triggers a premature response to starvation congruent with an apparently faster metabolism. Therefore, we speculate that selective pressure acts to maintain rDNA repeat number within an effective ‘optimal range’ that ensures rRNA levels can be readily tuned in response to environmental changes. More generally, we term this potential fitness defect ‘paralogous *cis*-rDNA interference’, whereby placement of two repeated and identical sequences in *cis* may cause interference with their function and ultimately lead to the loss of one locus.

These findings are noteworthy in the context of rDNA expansions observed in tumor cells, which are usually characterized by genome instability and rapid growth-associated metabolic states predominantly based on fast glucose uptake and fermentation (e.g., the Warburg effect).

## Methods

### Strains and culture conditions

All strains used in this study are listed in Extended Data Table 1 and they are derived from BY4741 and BY4742^55^ by transformation events. Deletion of the entire open reading frame of *SIR2* gene was performed using the CRISPR-Cas9 with a single gRNA and a ∼1 kb-long repair donor generated by fusion PCR on *SIR2*-flanking sequences. The thermosensitive mutation in *CDC15* (*cdc15-2* ts) was introduce with CRISPR-Cas9. Oligos used in this study are listed in Extended Data Table 4.

Strains were grown in rich medium (YPD: 1% bacto peptone (Difco), 1% bacto yeast extract (Difco) and 2% glucose; YPG: 1% bacto peptone (Difco), 1% bacto yeast extract (Difco) and 3% glycerol) or in synthetic complete medium (SC: 0.67% yeast nitrogen base without amino acids (Difco), supplemented with a mix of amino-acids and 2% glucose). Cells were grown at either 30°C or 24°C, the latter corresponds to the permissive temperature of the conditional thermosensitive mutant *cdc15-2* (see also methods in Lazar-Stefanita et al., 2022 in press).

### Serial dilution assay and growth curves

Yeast cultures from independent isolates of *n*=16 wild-type (WT) rDNA (BY4741), *n*=3 rDNA *CEN* distal (JL411), *n*=3 rDNA *CEN* proximal (JL665) and *n*=3 rDNA *CEN* proximal and distal (LS71) strains were grown in YPD liquid medium at 30°C to saturation. For serial dilution assay: yeast cultures in stationary phase were diluted to an optical density (OD) A_600_ = 0.1 and serially diluted (1:10) in sterile water then spotted on different media (YPD, YPG and SC) and incubated at different temperatures (30°C and 37°C) for 2 days. For growth curves: yeast cultures in stationary phase were diluted in fresh YPD medium to A_600_ = 0.05, 200 µl were transferred to 96 well plates and every minute the BioTek Eon microplate spectrophotometer was programmed to shake the plate and measure the A_600_ for a total of 24 h at 30°C. A_600_ values were imported in GraphPad Prism version 9 for Mac OS (GraphPad Software, San Diego, California USA, www.graphpad.com) and used to calculate mean and standard deviation for each isolate of each strain. Doubling time was calculated using the slope of the growth curves.

### rDNA Megablock engineering

A CRISPR-Cas9 based method was used to promote during the cut-and-paste rDNA locus relocation from the native location (flanked by chromosome 12 sequence) to an ectopic position generated by fusing telomere 15R and telomere 4L. The method was performed as described by Luo et al. *Nature* in 2018, except that two gRNAs cut the flanking regions of rDNA locus and the third gRNA cuts at the new ectopic insertion site. In addition, three donor templates were generated by fusion PCR, they include: (*i*) a donor template to rejoin the ends of the native locus upon rDNA deletion, (*ii*) a donor template for bridging the left flanking region of the rDNA locus to its new target location and (*iii*) a donor template for bridging the right flanking region of rDNA locus to its new target location. rDNA translocation was verified by PCR with primer pairs designed outside the newly formed chromosome junctions.

### Pulsed-field gel electrophoresis (PFGE) and Dual color Southern blot

Chromosomes from stationary yeast cell cultures were prepared in agar molds using the Certified Megabase Agarose (Bio-Rad CAT. 1613108) and pulsed-field gel electrophoresis (PFGE) was carried out as previously described, with running conditions recommended for either *S. cerevisiae* or *S. pombe* chromosomes. Specifically, megachromosomes were separated on a 0.8% agar gel in 1X TAE at 14°C. Run time was ∼72 h at 2V/cm with switch time of 20-30 min at an included angle of 106° ^16^. DNA size markers used were the *S. pombe* chromosomal DNA (Bio-Rad CAT. 170- 3633) and the *H. wingei* chromosomal DNA (Bio-Rad CAT. 170-3667).

Method for chromosome digestion: prior to the BamHI (NEB CAT. R0136L) incubation, agar plugs were equilibrated 1X Buffer3.1 (NEB) for 24 h at 4°C. BamHI digest was carried out for 24 h at 37°C in ∼500 µl final volume consisting of: ∼100 mg agar, 50 µl 10X restriction buffer and 10 µl BamHI enzyme (200 U). Agar molds treated with BamHI were then used for the PFGE and Southern blot.

The protocol for dual color Southern blot was optimized based on a previous study by Zavala et al.^56^. Following electrophoresis of BamHI digested agar plugs, the gel was washed in: depurination solution (0.25 M HCl) for 20 min, denaturation solution (0.5 M NaOH, 1.5 M NaCl) for 60 min and neutralization solution (0.5 M Tris, 1.5 M NaCl pH 7.5) for 30 min. Chromosomes were transferred by capillarity to a nylon membrane (Pall® 60208 Biodyne^TM^ B Membrane, Pore Size 0.45 µm CAT. 60208) using 10X SSC buffer (20X SSC: 1.5 M NaCl, 0.3 M Sodium citrate pH 7) for 72 h. After transfer the membrane was washed with 2X SSC for 5 min and baked at 80°C for 30 min. Prehybridization and hybridization steps were performed at 50°C in ULTRAhyb^TM^ Ultrasensitive Hybridization Buffer (Invitrogen Ambion CAT. AM8670). PCR products (∼1.1 kb-long flanking unique sequences on either side of each rDNA locus) were labelled using Klenow Fragment exo- (NEB CAT. M0212L) with either Biotin-16-dUTP (Roche CAT. 11093070910) or Digoxigenin-11-dUTP alkali-stable (Roche CAT. 11093088910) at 37°C for 1 h. Labelled DNA probes were cleaned up using Zymo-Spin I Columns (Zymo Research CAT. C1003-50), denatured at 95°C for 5 min and ∼300 ng of each were used for each hybridization experiment. The blot was washed twice with 2X SSPE buffer (20X SSPE Buffer: 0.02 M EDTA and 2.98 M NaCl in 0.2 M phosphate buffer pH 7.4; Sigma-Aldrich CAT. S2015) for 5 min at room temperature; twice with 2X SSPE, 1% SDS for 30 min at 55°C; twice with 0.1X SSPE for 15 min at 55°C. Blot was incubated in Blocking buffer Odyssey (LI-COR® CAT. 927-40000) in 1X Tris-Buffered Saline with 0.1% Tween 20 and 1% SDS for 3 h at room temperature.

The membrane was washed with 1X TBST for 5 min and incubated for 1 h at room temperature with rabbit anti-DIG antibody (working concentration 1:2500 in Blocking buffer Odyssey; ABfinity™ Rabbit Monoclonal, CAT. 700772). The primary antibody solution was washed out three times with 1X TBST for 15 min at room temperature. The membrane was incubated in the secondary antibody solution (working concentration 1:10000 in Blocking buffer Odyssey): Licor IRDye® 800CW Streptavidin, 0.5 mg (CAT. 926-32230) and LIRDye® 680RD Goat anti-Rabbit IgG (H + L), 0.5 mg (CAT. 926-68071) for 30 min at room temperature. Secondary antibody solution was discarded and the membrane was washed three times with 1X TBST for 15 min at room temperature before imaging acquisition using LI-COR Odyssey® Imager.

### Mating type switching

Here we used the same reagents and protocol described by Zhang et al.^42^.

### Sporulation and tetrad dissection

A detailed protocol for mating, sporulation and tetrad dissection of yeast with fused chromosomes was described by Luo et al.^16^.

### RNA extractions

Total RNA was extracted from 3 independent isolates of *n*=16 WT rDNA (BY4741), *n*=3 rDNA *CEN* distal (JL411), *n*=3 rDNA *CEN* proximal (JL665) and *n*=3 rDNA *CEN* proximal and distal (LS71) strains. Approximately 2×10^8^ cells were harvested from log phase cultures (1.5-2 x10^7^ cells/ml) grown in YPD medium at 30°C. Cell pellets were washed in RNase free water and resuspended in RNA lysis buffer (50 mM Tris-HCl pH 8, 100 mM NaCl). Cells were lysed mechanically using acid-washed glass beads (Sigma-Aldrich CAT. G8772-100G) at 4°C. The RNA was extracted by phenol:chloroform:isoamylalcohol (ThermoFisherScientific CAT. 15593) and ethanol precipitated. Extractions were treated with DNaseI (Agilent CAT. 600031) for 1 h at 37°C and RNA quality was verified by agarose gel in 1X TAE.

### RT-qPCR

The above total RNA extractions were serial diluted (1:10/100/1000) and used for RT-qPCR reactions with gene specific oligos (18S, 25S and TAF10). The reverse transcriptase (RT) reaction was performed according to the manufacturer protocol SuperScript™ IV Reverse Transcriptase (Invitrogen CAT. 18090050). Successively, quantitative PCR was performed using the LightCycler® 480 SYBR Green I Master (Roche CAT. 04887352001) and the standard amplification protocol with 45 cycles in a multi-well PCR plate 384. Ct values for each replicate were imported in GraphPad Prism version 9 for Mac OS (GraphPad Software, San Diego, California USA, www.graphpad.com) and used to calculate mean and standard deviation for each gene in each strain. The three plots generated, that correspond to each gene, show Ct variation in function of the RNA dilution factor and strain.

### RNA library preparation, sequencing and analysis

Stranded total RNA-seq libraries were prepared with the QIAseq stranded RNA library kit (Qiagen, CAT. 180743) following the manufacture’s recommendations using dual indexing primers. Additionally, the input total RNA was first processed with the QIAseq FastSelect rRNA Yeast kit to remove cytoplasmic and mitochondrial rRNA (Qiagen, CAT. 334215). Library quality was assessed using the ZAG 110 dsDNA kit on (Agilent, CAT. ZAG-110-5000) and run on the ZAG DNA analyzer system (Agilent). Libraries were then sequenced on an Illumina NextSeq 500 using the 75-cycle high output in paired end mode (Illumina, CAT. 20024906).

Differential gene analysis (Extended Data Table 2). RNA sequencing analysis was performed as previously described^57^ using kallisto/sleuth suite with minor modifications. Briefly, reads were processed by Illumina barcode and quality trimming with Trimmomatic^58^ and quality assessed with FastQC. Next, reads were aligned using the pseudo-alignment program kallisto^59^ to the reference S288C genome^60^, with annotations supplied with a gtf file containing all annotated RNAs in stranded paired end mode (options, --genomebam --gtf --fr-stranded). The output of kallisto was then used for data analysis in sleuth^61^. Volcano plots were generated using the Walds test false discovery rate adjusted *P* value (*p*<0.01) and log2 fold change (−2>Log2FC>2).

### Cell cycle synchronization

For both imaging and Hi-C experiments yeast cells, carrying either a single rDNA locus (LS88, LS95) or two rDNA loci in *cis* (LS90, LS103), were synchronized in either G1 with alpha-factor (Zymo Research, CAT. Y1001), or in late anaphase, using the thermosensitive mutant *cdc15-2*.

Method for G1 synchronization (LS88 and LS90): cells were grown to saturation in 5 ml SC medium at 30°C, the next day cells were subcultured in 150 ml fresh SC medium (optical density, OD, A600=0.5) for 2 h 30 min at 30°C then alpha-factor was added (final concentration: 25 µM) and cultures were incubated for additional 2 h 30 min at 24°C before imaging and formaldehyde crosslink.

Method for late anaphase synchronization (LS95 and LS103): cells were grown to saturation in 5 ml SC medium at 24°C, the next day they were subcultured in 150 ml fresh SC medium (A_600_=0.5) and grown for 5 h at 24°C then the temperature was switched to 37°C (non-permissive condition) and incubated for additional 3 h before imaging and formaldehyde crosslink.

The quality of G1 and anaphase synchronizations was visually monitored by microscopy hourly. For microscopy small aliquots (∼1-2 ml synchronized cultures) were spooned down at 3000g for 3 min and fixed in ethanol 70%; whereas, the remaining cultures were fixed with formaldehyde for Hi-C library preparation (see below).

### Protein tagging and imaging of the nucleolus

The cell cycle reorganization of the nucleolus within the nucleus was monitored using fluorescently tagged proteins at their endogenous C-terminus. Nuclear envelope was labelled with either mCherry (*NIC96::mCherry*) or mScarlet (*NUP49::mScarlet*), while the nucleolus with GFP (*NOP10::EGFP*) as described by Zhang et al.^42^ (see also strain genotype in Extended Data Table 1). Imaging was performed on the EVOS M7000 microscope using the Olympus X-APO 100 Oil, 1.45NA/WD 0.13mm (Oil) objective. Images were acquired as Z-stacks and visualized as max intensity projections using ImageJ^62^.

### Hi-C library preparation and data analysis

Hi-C libraries (Extended Data Table 3) were prepared from yeast cells harvested from either log phase or cell cycle synchronized cultures (see above protocols for cell synchronizations). Methods for yeast Hi-C library preparation and data analysis were performed as previously described by Lazar-Stefanita et al., 2017; 2022. Briefly, Hi-C reads were aligned using Bowtie 2^63^ and processed using command lines based on HiCLib tool developed in the Mirny lab (Imakaev et al., 2012). Normalized contact maps (Sequential Component Normalization)^64^ were 5 kb-binned and were plotted using MatLab R2018 (The MathWorks, Inc., Natick, Massachusetts, United States).

The Hi-C assay resulted crucial to validate the transplantation of the rDNA locus. We found that among the potential candidates - that were PCR positive for the rDNA transplantation - only one isolate had the entire locus relocated that resulted in the actual re-positioning of the rDNA boundary from the native location (centromere-distal: 2.75 Mb from the active centromeric sequence, *CEN15,* and 0.6 Mb from the nearest telomeric sequence, *TEL12R*) to the new ectopic position at the fusion junction between the left arm of former chromosome *15* and the right arm of former chromosome *4* (centromere-proximal: 0.75 Mb from *CEN15* and 2.52 Mb from *TEL12R*). Of note, in the isolates that were false positives by PCR for the rDNA megablock transplantation, the chromosome boundary formed by this locus remained at the original location (centromere-distal) as expected.

The Hi-C maps were used to evaluate the strength of each rDNA boundary in function of their distance from the centromere and locus number (single vs. double) across megachromosome *C*. For this analysis we firstly reasoned that the chromosome boundaries observed at the location of the rDNA loci on the 2D maps are ‘visual artifacts’ due to the combination of different structural characteristics at this locus. Indeed, the highly repetitive nature and the formation of a subnuclear compartment (nucleolus) appear to lower the contact frequency between the rDNA-flanking regions. Along the same lines we also expected that differences in structure and/or size between rDNA loci would have an effect on the strength of the correspondent chromosome boundaries. In other words, bigger the rDNA locus (‘bulkier’ the nucleolus) lower the contact frequency between its flanking sequences, therefore stronger the boundary. To compare between the *CEN* proximal and *CEN* distal rDNA boundaries, located in single or double copy on megachromosome *C*, we used normalized maps (5 kb-binned) to quantify contacts between 500 kb-long regions (flanking either the right or the left end of each locus) and the 1.9 Mb region upstream/downstream or enclosed by two loci (see highlighted regions in blue on chr *C* maps in Figure 3a). Spread point plots, where each point corresponds to total contacts made by each 5 kb bin, were generated for each rDNA boundary using Matlab. *P* values were calculated using the Kolmogorov–Smirnov test (K–S test) function in MATLAB 2018a (The MathWorks, Inc., Natick, Massachusetts, United States).

Log2 ratios of Hi-C contact maps were used for their pairwise comparison and the contact probability decay in function of the genomic distance, *p*(*s*), was computed to assess the status of the chromatin fiber. Both these analyses were performed as previously described by Lazar-Stefanita et al., 2017; 2022 in press.

## Acknowledgments

We acknowledge all members of Boeke lab for their assistance. We thank Aleksandra Wudzinska and Gregory W. Goldberg for their assistance and helpful comments on the manuscript. This work was supported by National Science Foundation grants MCB-1616111 and MCB-1921641 and NIH grant RM1HG009491 to JDB.

## Author Contributions

LLS and JDB designed the research project. LLS performed the experiments with help from: JL that relocated the rDNA array, WZ that built *n*=16 with rDNA on heterologous chromosomes and MH that performed and analyzed RNAseq. LLS wrote the manuscript and all authors contributed to editing.

## Additional Information

Genome-wide and RNA sequencing files were deposited at the NCBI database as FASTQ files. See BioProject accession number PRJNA847233 for the SRA data. The reviewer link: https://dataview.ncbi.nlm.nih.gov/object/PRJNA847233?reviewer=6h4lqpu2o2epg0f4ta40dlmqo 4

## Extended Data Tables

**Extended Data Table 1.** List of strains used in this study.

| Strain name | Parent | Genotype | No. Chr. | rDNA location | Ref. |
| --- | --- | --- | --- | --- | --- |
| BY4741 | S288C | *MATa his3Δ1 leu2Δ0 met15Δ0 ura3Δ0* | 16 | Chr 12 | (Brachmann et al., 1998) |
| WZ456 | BY4742 | *MATa his3Δ1 leu2Δ0 lys2Δ0 ura3Δ0 rdn::C-S.ITS-LEU2 NOP10::EGFP-KanMX, NIC96::mCherry-HIS3* | 16 | Chr 12 | (Zhang et al., 2017) |
| WZ457 | BY4742 | *MATa his3Δ1 leu2Δ0 lys2Δ0 ura3Δ0 rdn::NatMX4 NOP10::EGFP-KanMX, NIC96::mCherry-HIS3 YCR018C / C-S.ITS-LEU2 / YCR019W* | 16 | Chr 3 | (Zhang et al., 2017) |
| WZ499 | Spore of yWZ456 X yWZ457 | *MATa his3Δ1 leu2Δ0 lys2Δ0 ura3Δ0 rdn::C-S.ITS-LEU2 NOP10::EGFP-KanMX, NIC96::mCherry-HIS3, YCR018C / C-S.ITS-LEU2 / YCR019W* | 16 | Chr 12 + Chr 3 | (Zhang et al., 2017) |
| JL411 | BY4741 | *MATa his3Δ1 leu2Δ0 met15Δ0 ura3Δ0* | 3 | Chr C distal | (Luo et al., 2018a) |
| JL665 | JL411 | *MATa his3Δ1 leu2Δ0 met15Δ0 ura3Δ0* | 3 | Chr C proximal | This work |
| LS88 | JL665 | *MATa his3Δ1 leu2Δ0 met15Δ0 ura3Δ0 NOP10::EGFP-KanMX NUP49::mScarlet-S.p.HIS5* | 3 | Chr C proximal | This work |
| LS95 | LS88 | *MATa his3Δ1 leu2Δ0 met15Δ0 ura3Δ0 cdc15-2(Ts) NOP10::EGFP-KanMX NUP49::mScarlet-S.p.HIS5* | 3 | Chr C proximal | This work |
| LS70 | JL665 X JL411 | *MATa/alpha his3Δ1/ his3Δ1 leu2Δ0/ leu2Δ0 met15Δ0/met15Δ0 ura3Δ0/ura3Δ0* | 3 | Chr C proximal & Chr C distal | This work |
| LS71-C1&C2 | Spore of LS70 | *MATalpha his3Δ1 leu2Δ0 met15Δ0 ura3Δ0* | 3 | Chr C proximal + distal | This work |
| LS71-C3 | Spore of LS70 | *MATa his3Δ1 leu2Δ0 met15Δ0 ura3Δ0* | 3 | Chr C proximal + distal | This work |
| LS90 | LS71-C3 | *MATa his3Δ1 leu2Δ0 met15Δ0 ura3Δ0 NOP10::EGFP-KanMX NUP49::mScarlet-S.p.HIS5* | 3 | Chr C proximal + distal | This work |
| LS91 | Haploids of JL411 | *MATa/alpha his3Δ1/ his3Δ1 leu2Δ0/ leu2Δ0 met15Δ0/met15Δ0 ura3Δ0/ura3Δ0* | 3 | Chr C distal & Chr C distal | This work |
| LS92 | Haploids<br>of JL665 | *MATa/alpha his3Δ1/ his3Δ1 leu2Δ0/*<br>*leu2Δ0 met15Δ0/met15Δ0 ura3Δ0/ura3Δ0* | 3 | Chr C<br>proximal &<br>Chr C<br>proximal | This work |
| LS93 | LS71-C1<br>X<br>LS71-C3 | *MATa/alpha his3Δ1/ his3Δ1 leu2Δ0/*<br>*leu2Δ0 met15Δ0/met15Δ0 ura3Δ0/ura3Δ0* | 3 | Chr C<br>proximal +<br>distal &<br>Chr C<br>proximal +<br>distal | This work |
| LS103 | LS90 | *MATa his3Δ1 leu2Δ0 met15Δ0 ura3Δ0*<br>*cdc15-2(Ts) NOP10::EGFP-KanMX*<br>*NUP49::mScarlet-S.p.HIS5* | 3 | Chr C<br>proximal &<br>distal | This work |
| LS130<br>(LS131) | JL411 | *MATa his3Δ1 leu2Δ0 met15Δ0 ura3Δ0*<br>*sir2Δ* | 3 | Chr C distal | This work |
| LS141<br>(LS142) | Diploid<br>(*SIR2/*<br>*sir2Δ*) | *MATa or alpha his3Δ1 leu2Δ0 met15Δ0*<br>*ura3Δ0 sir2Δ* | 3 | Chr C<br>proximal &<br>distal | This work |
| LS143<br>(LS144) | Diploid<br>(*sir2Δ/*<br>*sir2Δ*) | *MATa or alpha his3Δ1 leu2Δ0 met15Δ0*<br>*ura3Δ0 sir2Δ* | 3 | Chr C<br>proximal &<br>distal | This work |

**Extended Data Table 2 RNA seq. list of differentially expressed genes.**

Data presented as excel spreadsheet file.

**Extended Data Table 3.** Hi-C libraries. For each library it reports: number of chromosomes per strain (*n*), number of rDNA per chromosome, synchronization method and total Hi-C contacts.

| Strain name | Karyotype<br>(*n*, #rDNA-position) | Synchronization<br>method | Total<br>paired-end<br>reads | Aligned<br>paired-end<br>reads | Total contacts<br>in 5 kb-binned<br>maps |
| --- | --- | --- | --- | --- | --- |
| JL411 | 3, 1-distal | none | 24770805 | 19642950 | 5273769 |
| JL665 | 3, 1-proximal | none | 24447393 | 19581599 | 3662044 |
| LS70 | 3, 1-distal + 1-prox | none | 34383730 | 27174746 | 4918384 |
| LS71-C1 | 3, 1-distal + 1-prox | none | 29910065 | 20997502 | 3292606 |
| LS71-C2 | 3, 1-distal + 1-prox | none | 26807590 | 18886934 | 3585706 |
| LS71-C3 | 3, 1-distal + 1-prox | none | 32026043 | 23014301 | 4020070 |
| LS88 | 3, 1-proximal | G1 | 12605198 | 8902009 | 5362318 |
| LS90 | 3, 1-distal + 1-prox | G1 | 8820734 | 5893335 | 4041830 |
| LS95 | 3, 1-proximal | Anaphase *cdc15-2* | 12933900 | 9479614 | 6106239 |
| LS103-C1 | 3, 1-distal + 1-prox | Anaphase *cdc15-2* | 15026254 | 10109053 | 3904083 |
| LS103-C2 | 3, 1-distal + 1-prox | Anaphase *cdc15-2* | 28339485 | 20719596 | 4170357 |

**Extended Data Table 4 List of primers used in this study.**

Data presented as excel spreadsheet file.

## Extended Data Figures and Figure Legends

**Extended Data Figure 1.**
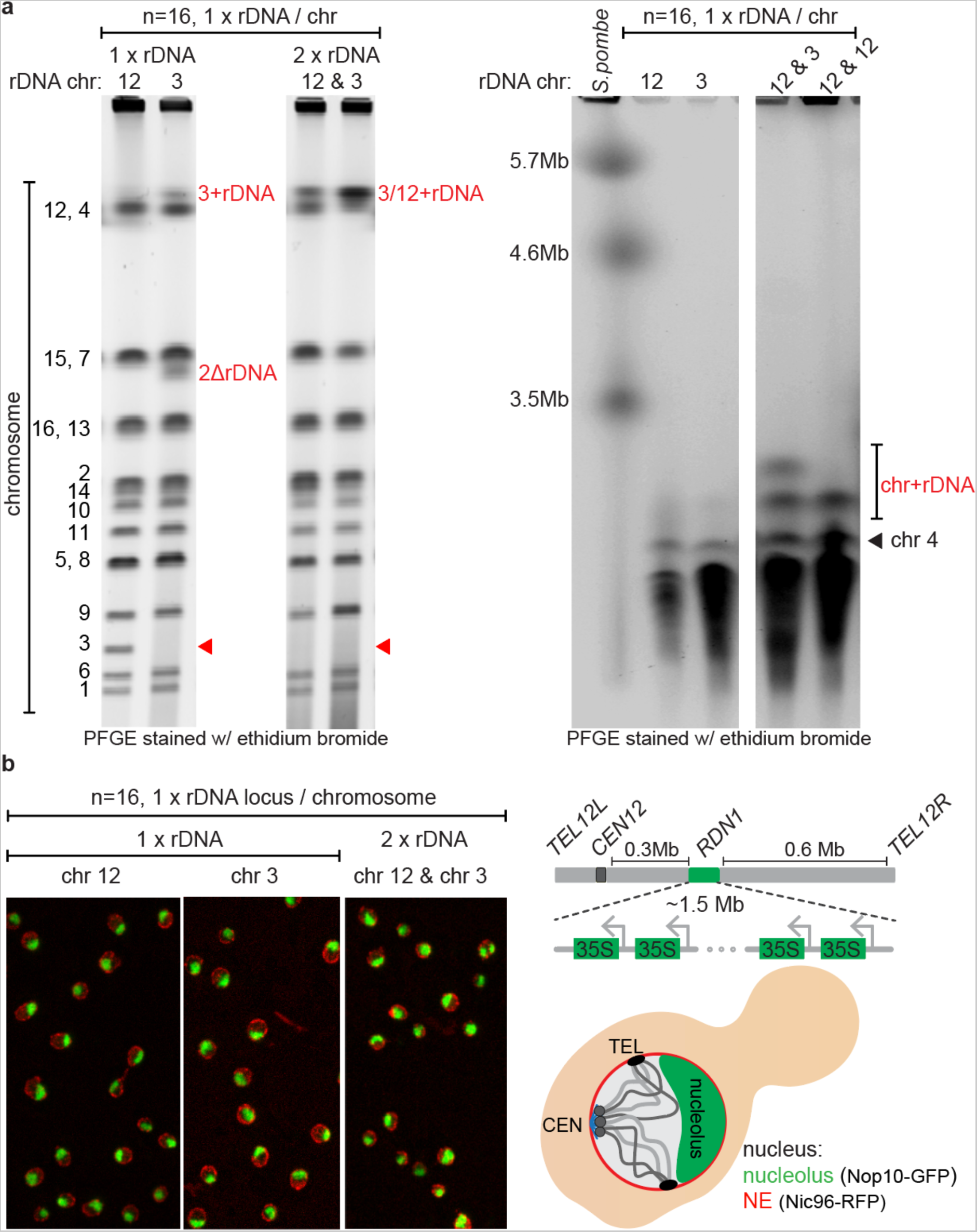
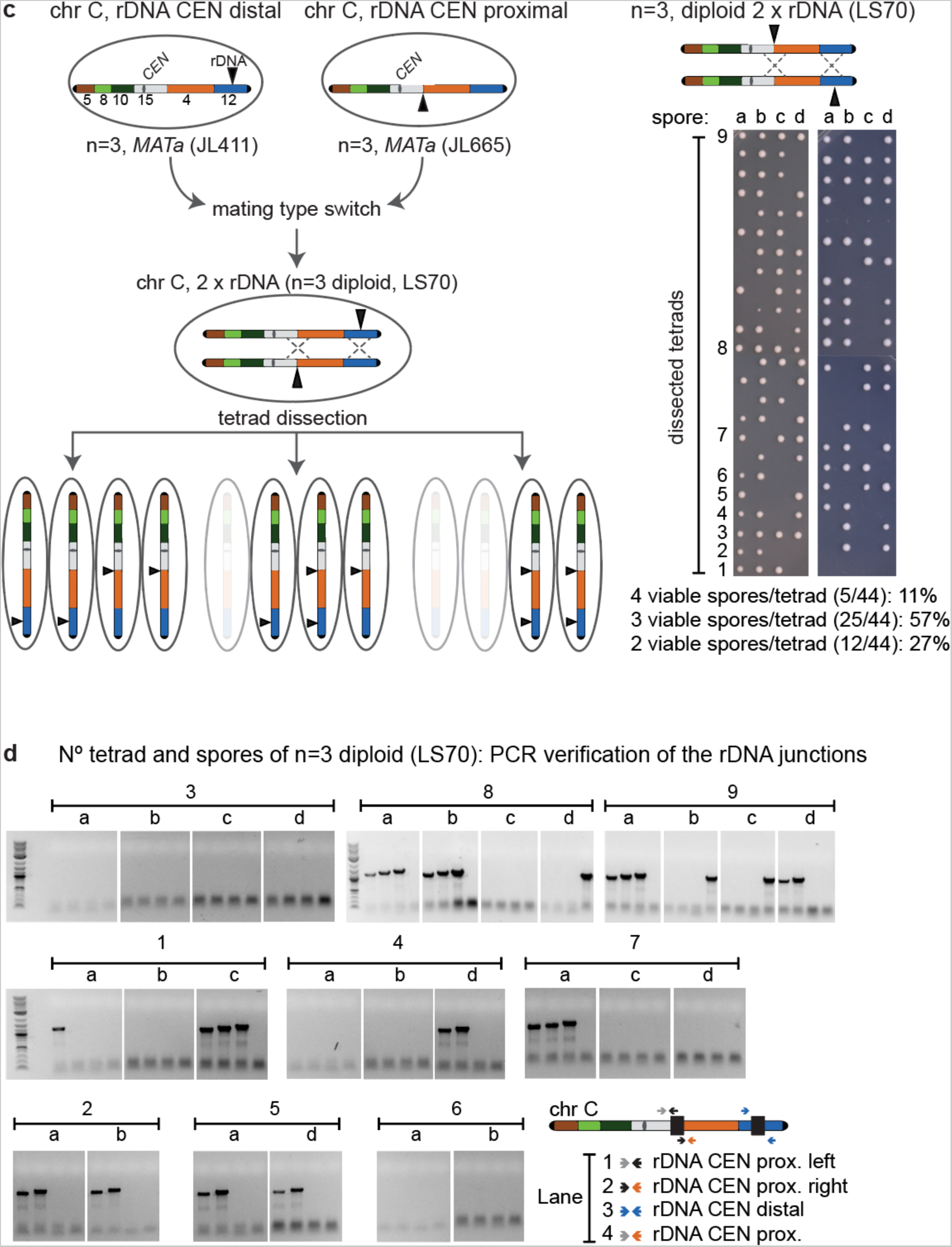
rDNA locus transplantation and duplication at ectopic chromosomal locations. **a** Chromosome separation by PFGE of wild-type yeasts with 16 chromosomes (*n*=16) with single or two rDNA loci on different chromosomes (red indicates chromosomes that have lost or gain the rDNA locus). Two distinct programs are used for chromosome separation: *S. cerevisiae* on the left and *S. pombe* on the right. **b** Representative microscopy images of yeast nuclei with single or two rDNA loci as described in Figure 1a, b. **c** Diagram showing the strategy used to duplicate the rDNA locus within the same arm of megachromosome *C* in *n*=3 (chr *C* is colorcoded according to the chromosome fusion design). Right panels show spores of individual tetrads (asci) of *n*=3 diploid with two rDNA loci in asymmetric *trans* locations. **d** Verification of rDNA duplication by junction PCR, sampling 4 different sequences in 27 spores originated from 9 different tetrads. Positive spores for rDNA duplication show PCR products in lane 1 and 2 only.

**Extended Data Figure 2.**
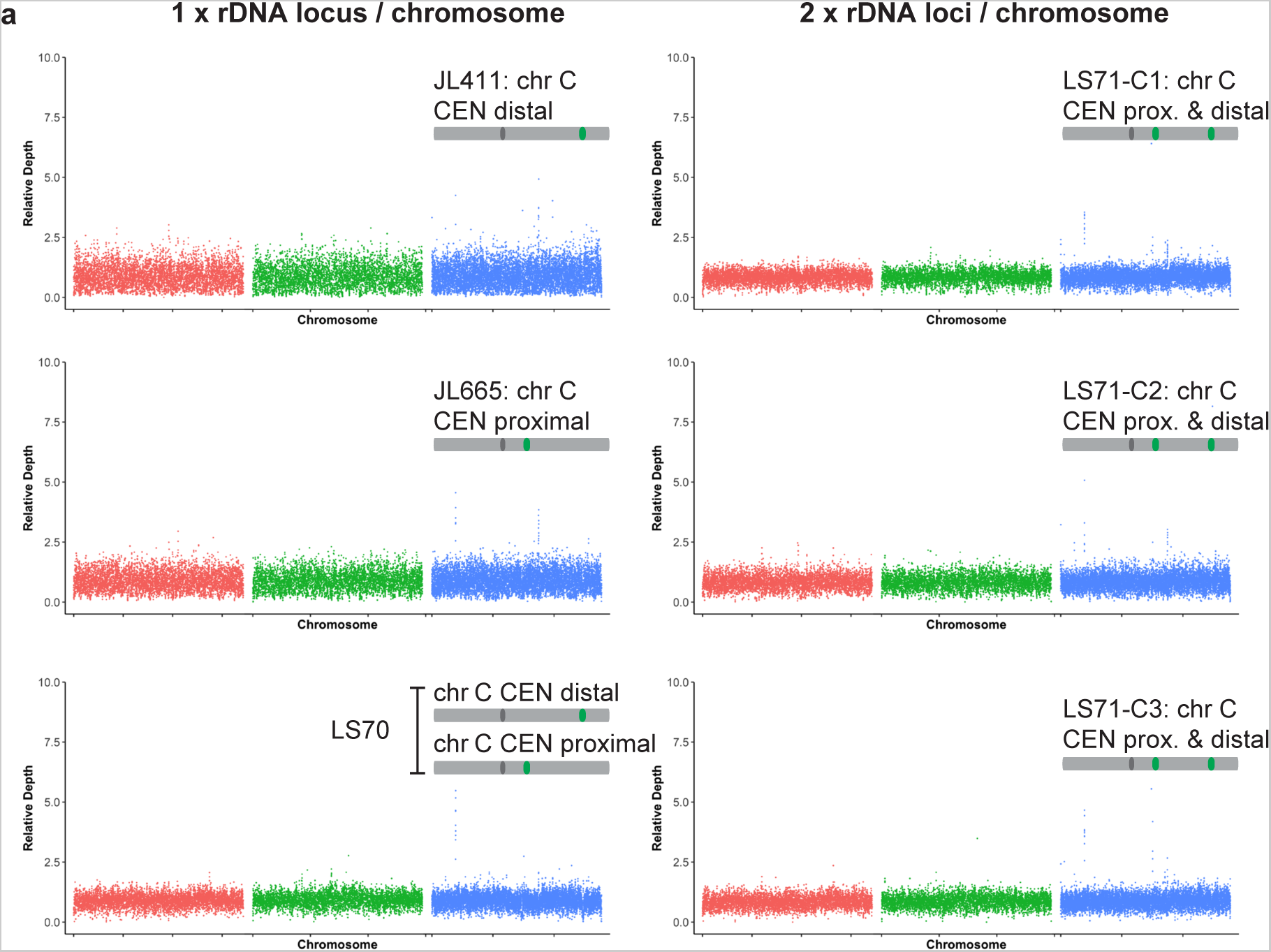
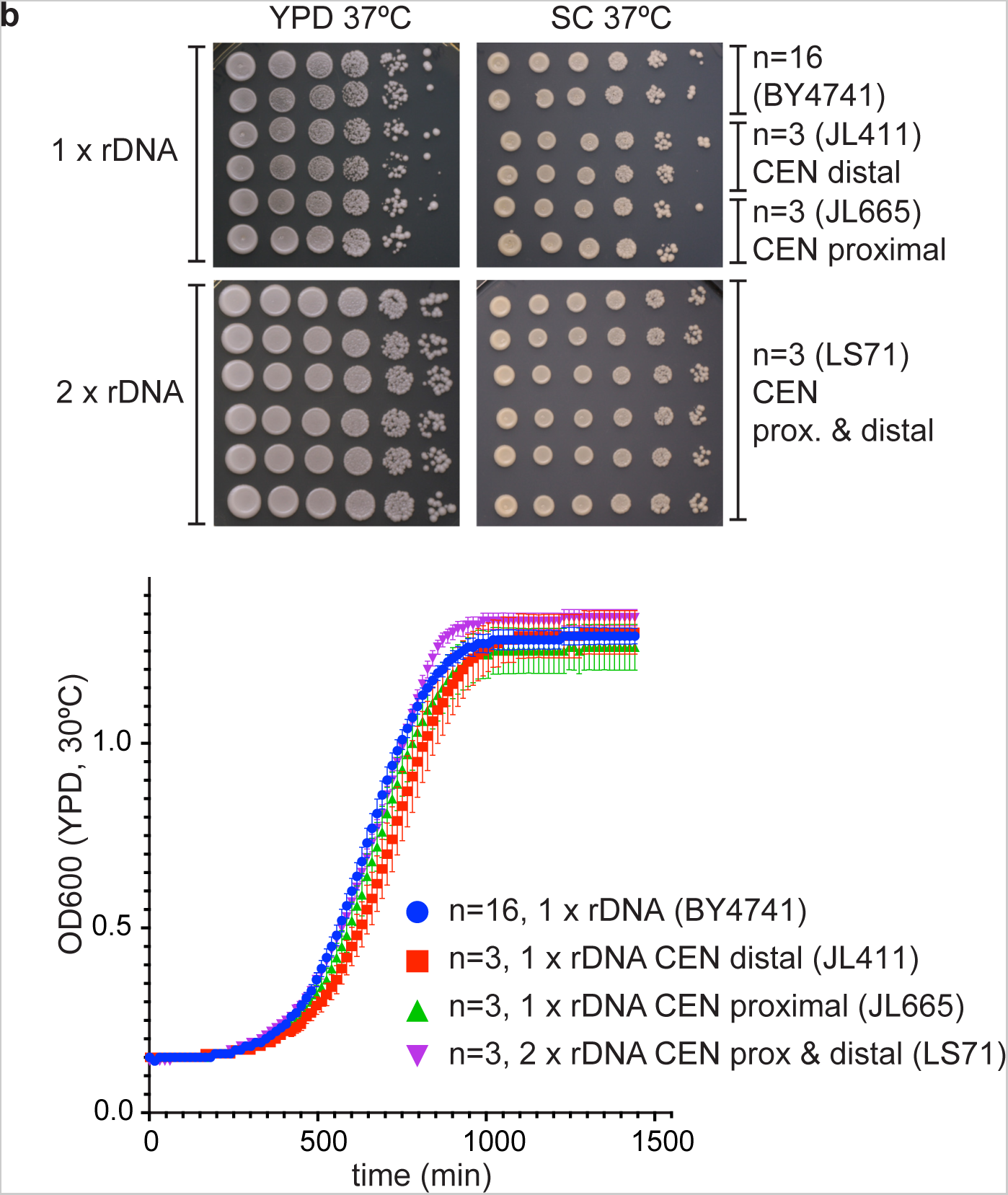
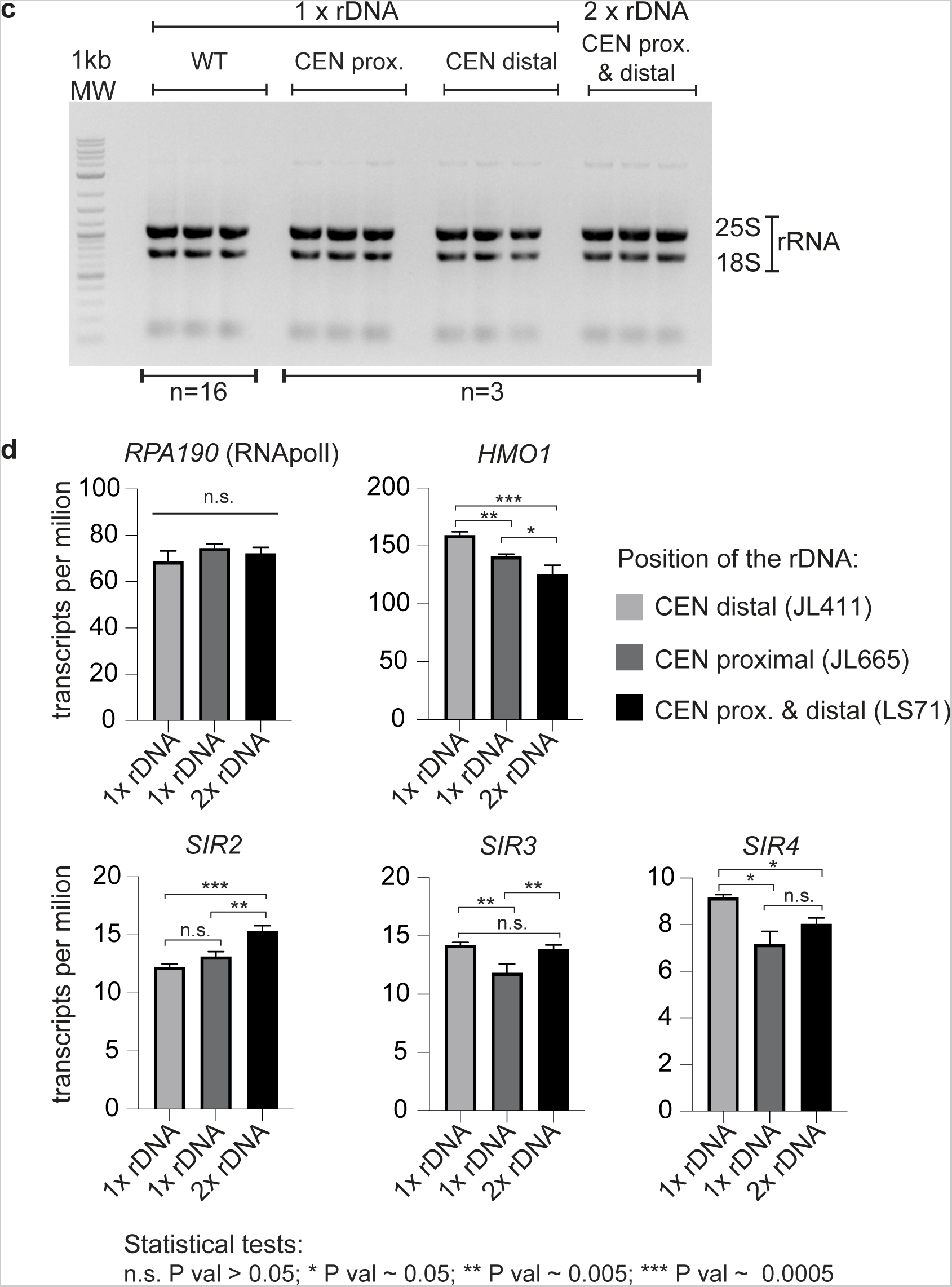
Effects of rDNA duplication on genome stability and cell fitness. **a** Relative genome sequencing depth plots (ggplot2 function in R) of *n*=3 strains with a single rDNA locus on chr *C* (haploids: JL411 and JL665 with either *CEN* distal or *CEN* proximal locus, respectively; diploid: LS70 with *CEN* distal and proximal) or two loci on chr *C* (independent haploids: LS71 C1 to C3). Sequencing depth for each megachromosome is colorcoded differently; chr *C* is displayed on the top right of each plot. **b** Fitness assays of *n*=16 and *n*=3 isolates. Top panels: serial dilution assays on different growing conditions at 37°C see also Figure 2b. Bottom panel: growth curves reporting optical densities (OD^A600^) in YPD at 30°C. **c** Agarose gel reporting the quality of the total RNA extracted from *n*=16 and *n*=3 isolates with different rDNA number and position relative to Figure 2c. **d** Bar plots showing transcriptional changes of a specific subset of genes (mainly involved in the metabolism of ribosomal DNA and RNA) in *n*=3 with a different number and position of the rDNA locus see also Figure 2d.

**Extended Data Figure 3.**
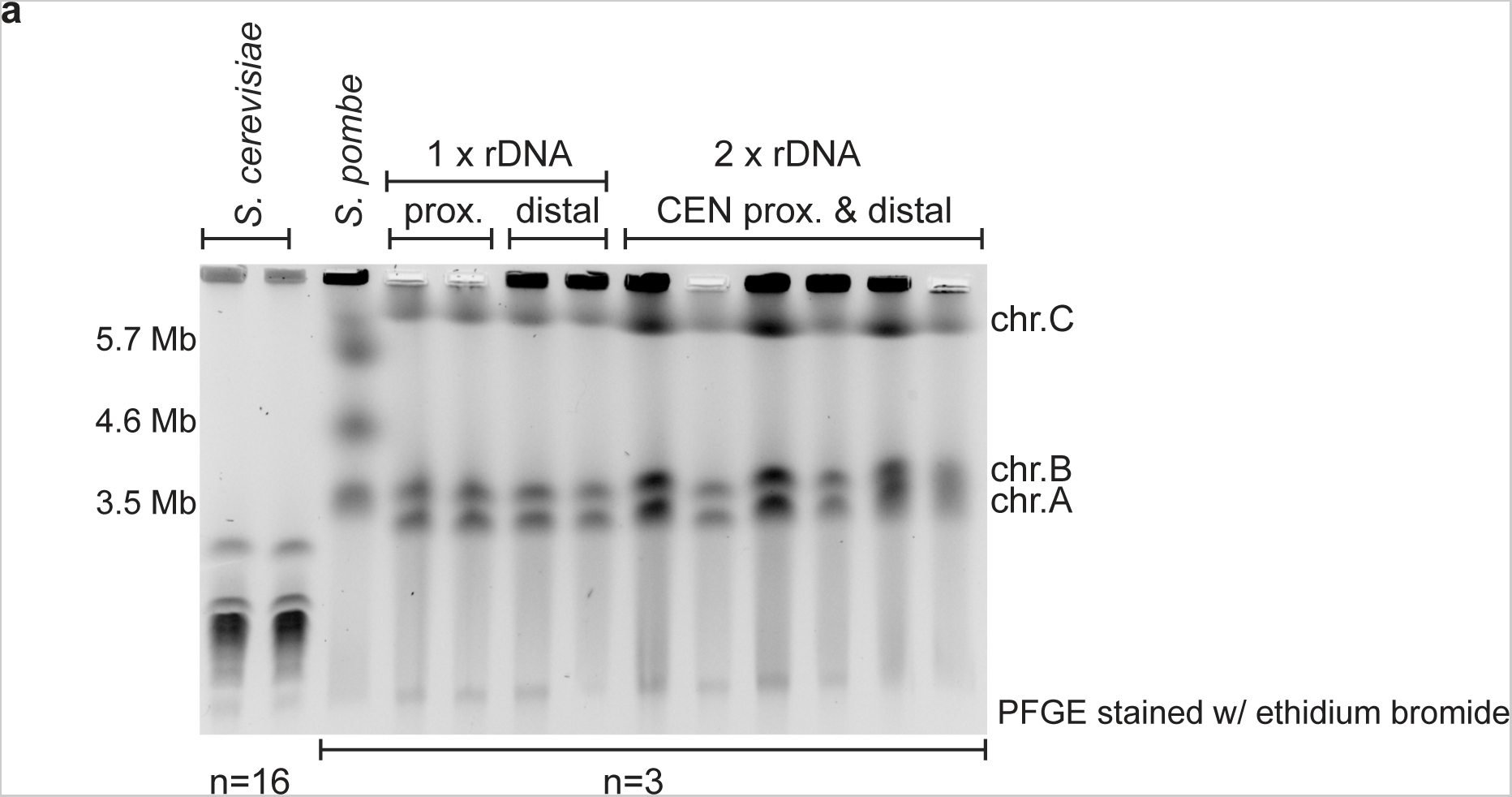
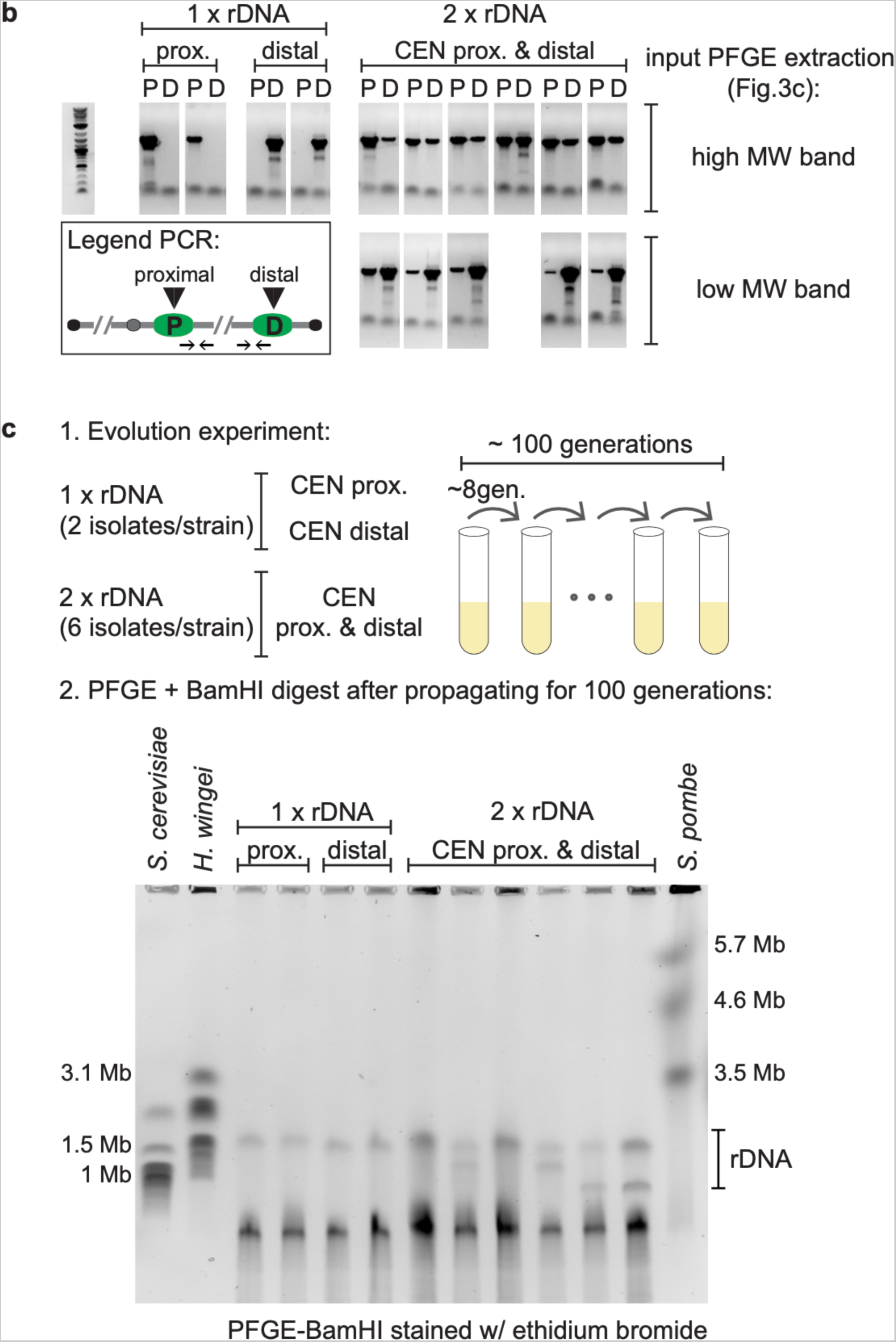
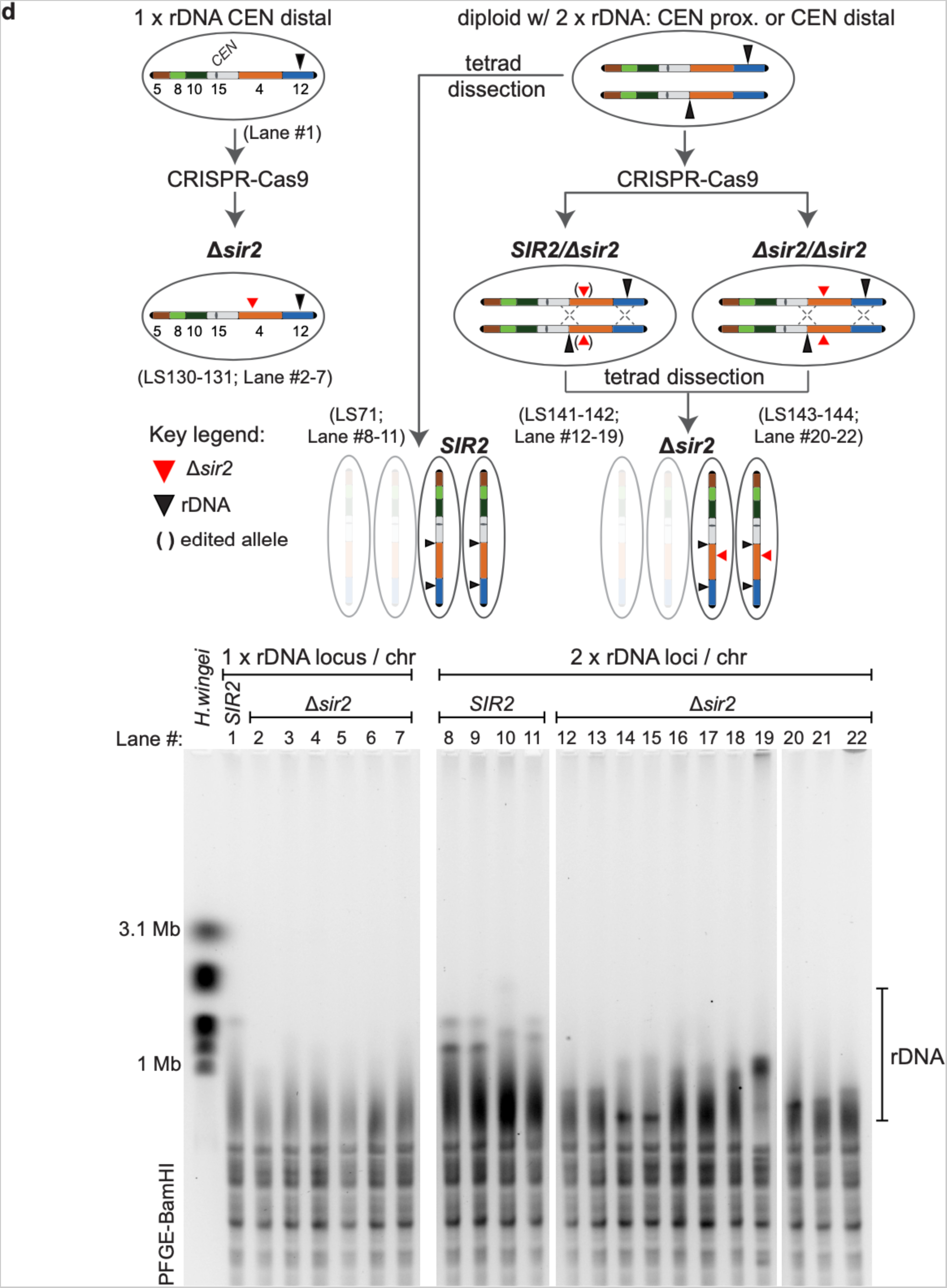
Stability of the rDNA locus in function of chromosome position and silencing. **a** Chromosome separation by PFGE of different isolates of *n*=16 and *n*=3 strains, with either one or two rDNA loci. PFGE run specifications: *S. pombe* program for mega-size chromosome separation. **b** PCR-validations of rDNA proximal (P) and distal (D) junctions. DNA template extracted from PFGE, rDNA bands that migrated at high or low molecular weights in Figure 3c. **c** Schematics of the evolution experiment in *n*=3 strains with a different number and position of the rDNA locus. Cell populations from different isolates were evolved long-term (∼100 generations) in batch cultures and used to assess the size of the rDNA arrays using PFGE and BamHI digest. PFGE program as described in panel (a). **d** Diagram of the approach used to assess rDNA array stability in function of Sir2. CRISPR-Cas9 was used to delete *SIR2* gene in *n*=3 haploid with one rDNA locus, and in the diploid with two rDNA loci in *trans* and asymmetric. Haploid spores with two *cis* rDNA loci obtained from sporulation of different diploids (*SIR2/SIR2, SIR2/sir2, sir2/sir2*) were used to assess stability of the rDNA arrays by PFGE and BamHI digest. PFGE program as described in panel (a).

**Extended Data Figure 4.**
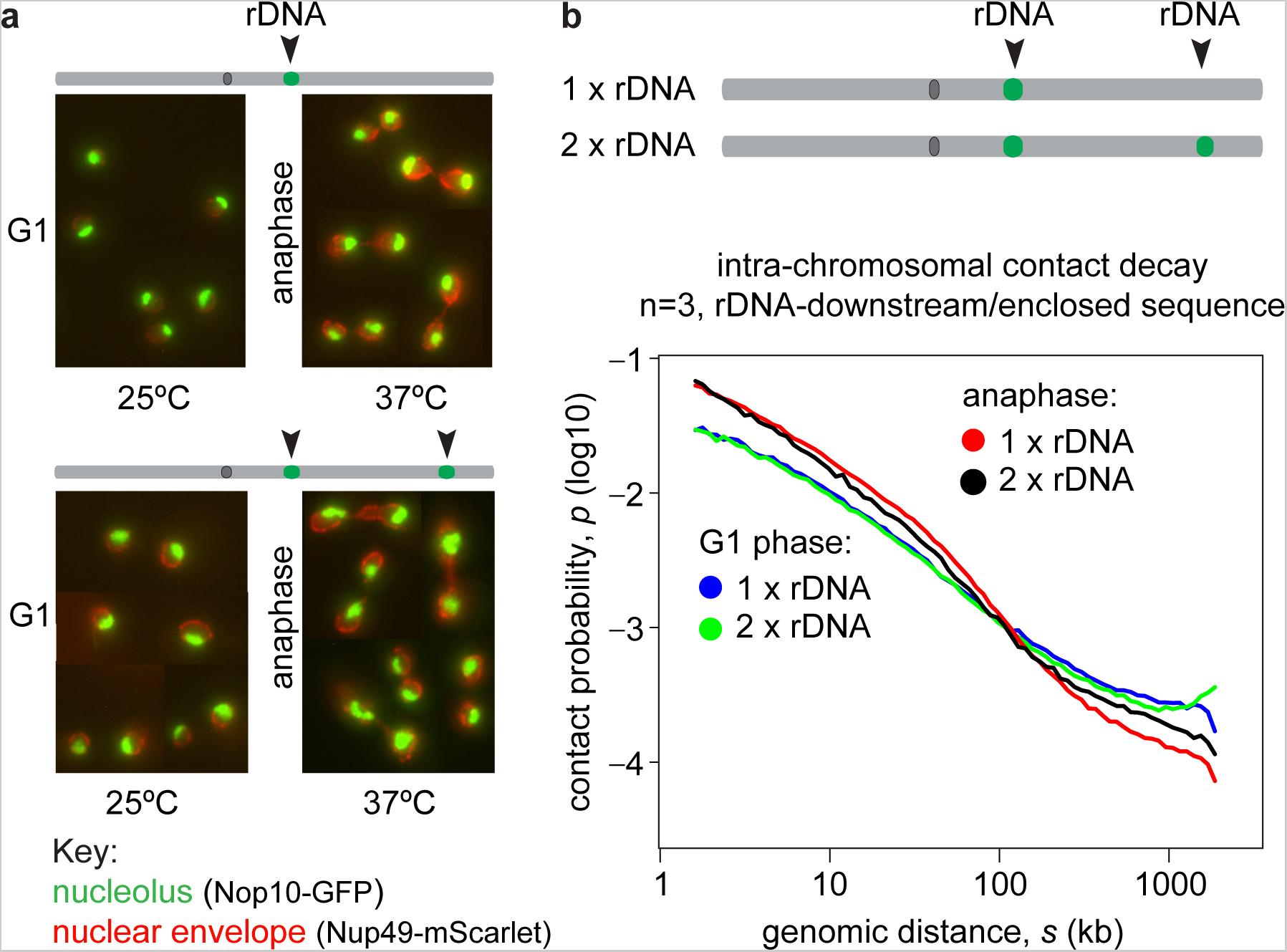
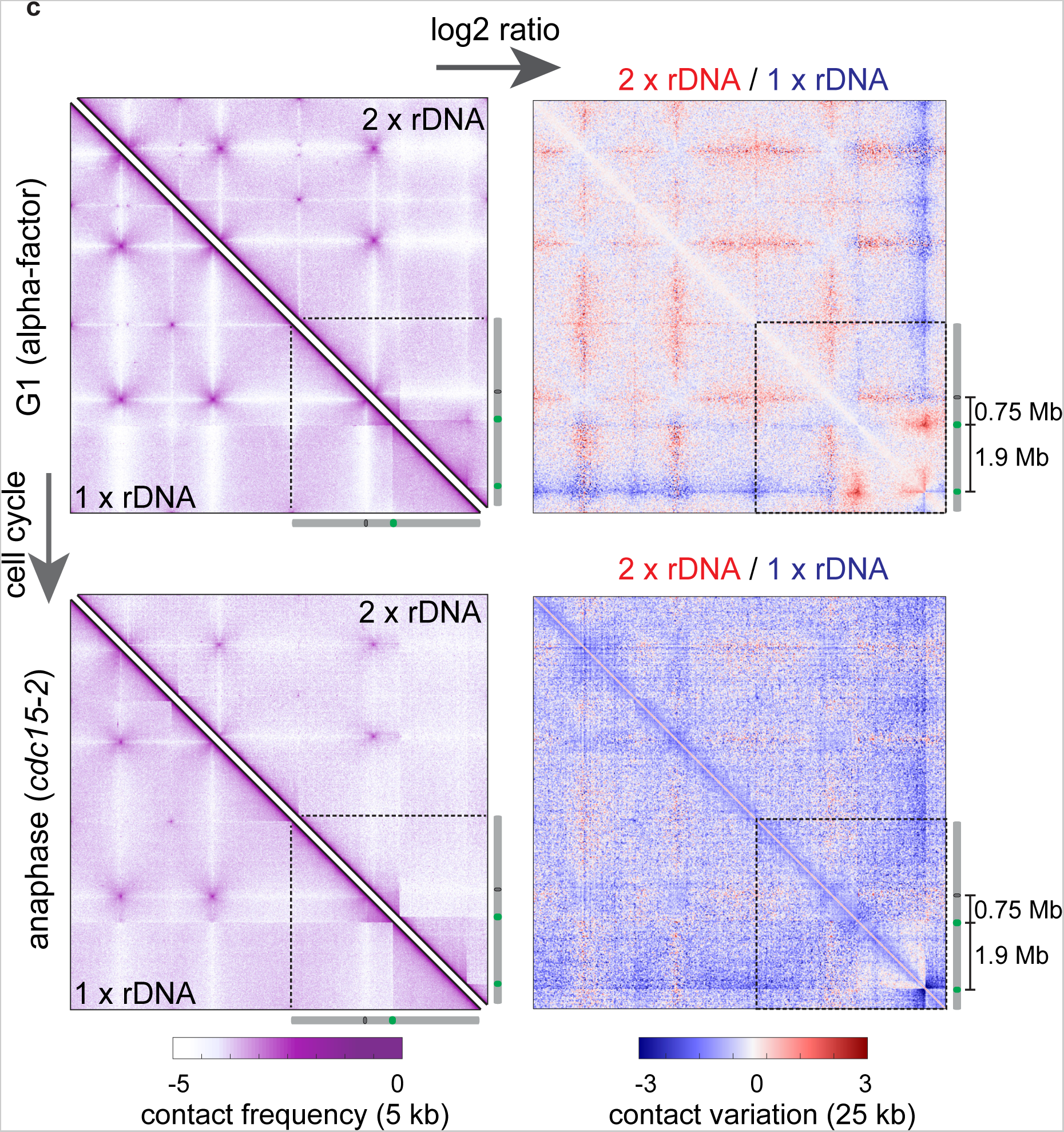
Cell cycle reorganization of the chromosome with twin rDNA loci in *cis*. **a** Representative microscopy images of *n*=3 nuclei, with one or two rDNA loci, in cells synchronized in G1 (alpha-factor, growth at 25°C) and late anaphase (*cdc15-2* ts, growth at 37°C). **b** Contact probability decay *p*(*s*) in G1 vs. anaphase: shows the average decay of the intrachromosomal contact frequency p between loci with respect to their genomic distance *s*. **c** Left panels: Hi-C maps of megachromosomes in *n*=3, with one or two rDNA on chr *C* (dashed outline), from cells synchronized in G1 (top triangle maps) and late anaphase (bottom triangle maps). Right panels: contact variation (log2-ratio) of the maps shown on the left (G1: top square map and anaphase: bottom square map).

**Extended Data Figure 5.**
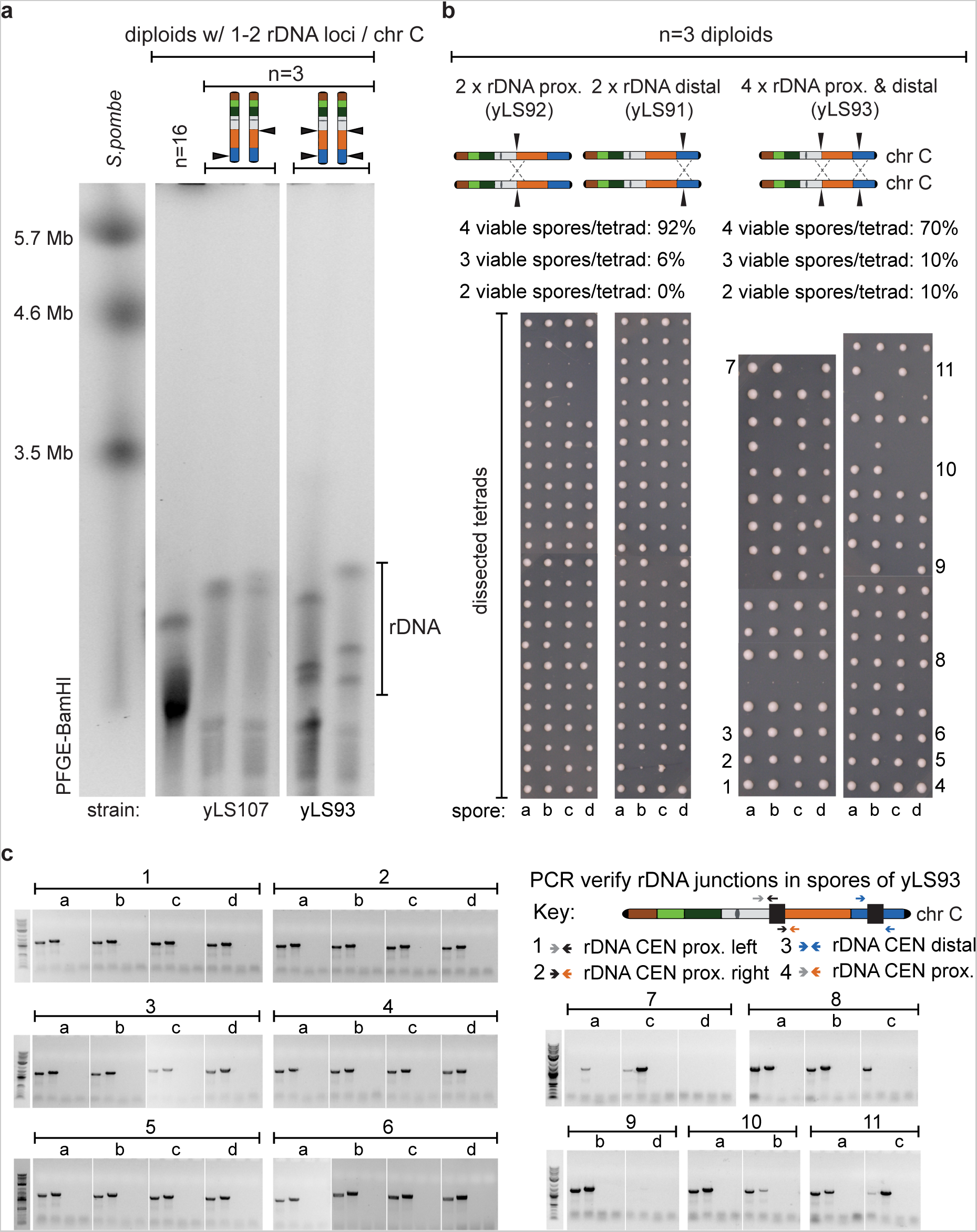
Duplicated rDNA loci may lead to chromosome mispairing in meiosis. **a** PFGE and BamHI digest of diploids of *n*=16 and *n*=3 strains, with different number and position of the rDNA locus. Run specifications: *S. pombe* program for mega-size chromosome separation. **b** Sporulation experiments of *n*=3 diploids with single and double pairs of rDNA loci. Percentage of tetrads (asci) with 2, 3 or 4 viable spores is indicated for each strain. **c** PCR verification of rDNA junctions, sampling 4 different sequences in 36 spores originated from 11 different tetrads.

